# Single-cell antigen-specific activation landscape of CAR T infusion product identifies determinants of CD19 positive relapse in patients with ALL

**DOI:** 10.1101/2021.04.15.440005

**Authors:** Zhiliang Bai, Steven Woodhouse, Dongjoo Kim, Stefan Lundh, Hongxing Sun, Yanxiang Deng, Yang Xiao, David M. Barrett, Regina M. Myers, Stephan A. Grupp, Carl H. June, J. Joseph Melenhorst, Pablo G. Camara, Rong Fan

**Author notes:** Correspondence to: J.J.M., P.G.C., R.F. These authors contributed equally to this work (Zhiliang Bai, Steven Woodhouse). These authors jointly directed this work.

## Abstract

Chimeric antigen receptor-modified (CAR) T cells targeting CD19 have mediated dramatic responses in relapsed/refractory acute lymphoblastic leukemia (ALL), yet a notable number of patients have CD19-positive relapse within one year of treatment. It remains unclear if the long-term response is associated with the characteristics of CAR T cells in infusion products, hindering the identification of biomarkers to predict therapeutic outcomes prior to treatment. Herein we present 101,326 single cell transcriptomes and surface protein landscape from the CAR T infusion products of 12 pediatric ALL patients upon CAR antigen-specific stimulation in comparison with TCR-mediated activation and controls. We observed substantial heterogeneity in the antigen-specific activation states, among which a deficiency of Th2 function was associated with CD19-positive relapsed patients (median remission 9.6 months) compared with very durable responders (remission>54 months). Proteomic profiles also revealed that the frequency of early memory T cell subsets, rather than activation or co-inhibitory signatures could distinguish CD19-positive relapse. Additionally, a deficit of type 1 helper and cytotoxic effector function and an enrichment for terminally differentiated CD8+ T cells exhibiting low cytokine polyfunctionality was associated with initial non-responders. By contrast, the single-cell transcriptomic data of unstimulated or TCR-activated CAR T cells failed to predict clinical responses. In aggregate, our results dissect the landscape of CAR-specific activation states in infusion products that can identify patients who do not develop a durable response to the therapy, and unveil the molecular mechanisms that may inform strategies to boost specific T cell function to maintain long term remission.

## Introduction

Children with chemotherapy-resistant and/or refractory (r/r) acute lymphoblastic leukemia (ALL) have dismal prognosis (*1, 2*). Emerging as a potential immunotherapeutic approach, autologous chimeric antigen receptor (CAR) T cellular therapy targeting the B-cell surface protein CD19 (CART19) has produced remarkable outcomes in the treatment of B cell malignancies (*3-7*). Multicenter clinical trials have demonstrated that 70-90% of patients with B-ALL achieve complete remission after receiving CART19 products (*6-9*), providing a potentially curative option for these patients. However, the remissions in a significant fraction of subjects are short-lived and 30-60% of treated patients relapse within one year (*8-11*), let alone ∼20% of ALL patients and over half of patients with large B cell lymphoma (LBCL) or chronic lymphocytic leukemia (CLL) fail to enter initial remission after CART19 infusion (*12-15*).

These disparities in therapeutic efficacy have evoked the efforts to interrogate the molecular determinants of long-term remission. It has been well accepted that robust *in vivo* CAR T cell engraftment and expansion is a prerequisite for inducing antitumor efficacy (*6, 16*), which is impacted by several CAR T product characteristics such as CAR construct design (*16, 17*) and early memory T cell frequencies (*18-20*). By characterizing the pre-manufacture starting T cell phenotype, an elevated frequency of CD27+CD45RO-CD8+ T cells (*21*), a higher percentage of naive and early memory T cell composition (*22*) and a decreased frequency of LAG-3+/TNF-α^lo^CD8+ T cells (*23*) were found to be associated with sustained remission. Additionally, tumor-intrinsic signatures, including the impaired death receptor signaling in tumor cells (*24*) and a high pretreatment disease burden (*8*), also contributed to observed therapeutic failure and reduced remission duration.

However, the molecular mechanisms that underlie the acquired resistance to the CAR T treatment are still unclear. Although antigen loss could explain a majority of CD19-negative disease relapse after receiving CART19 therapy (*25-27*), the mechanisms of CD19-positive relapse remain elusive. We hypothesized that the functional capacity of CAR T cells in the infusion product could be an essential factor determining long-term therapeutic response. By combining single cell RNA sequencing (scRNA-seq) and co-measurement of a panel of surface proteins (CITE-seq) (*28*), we have investigated the single-cell activation landscape of CAR T infusion products from 12 pediatric r/r ALL patients upon CAR-specific stimulation or TCR-mediated activation, in comparison with unstimulated controls. We find that CD19-positive relapsed patients (*n* = 5) have a deficit of CAR T cells capable of inducing a T helper type-2 (Th2) functional response and of maintaining stem cell-like memory and central memory states compared with very durable responders (*n* = 5), whereas the expression levels of activation- or co-inhibitory-related proteomic markers are comparable between two groups. Moreover, we find that patients with no initial response (*n* = 2) are unable to mount a strong helper and cytotoxic type-1 (Th1/Tc1) response, with a deficit of cells in this state and an enrichment of terminally differentiated CD8+ cells. These results are only manifest upon antigen-specific stimulation. In contrast, unstimulated or TCR-mediated activation states fail to predict clinical responses. These findings provide new insights into the molecular characteristics that underlie CD19-positive relapse via single-cell multi-omics profiling of CAR T activation states and computational dissection of functional modules and biological pathways associated with clinical outcomes.

## Results

### Single cell multi-omics profiling of CD19 CAR T infusion products from ALL patients

We analyzed the infusion products of 12 pediatric and young adult patients (median age 11.15 years; range 4.9-21.5 years) from a phase I/IIA CART19 clinical trial of r/r B-ALL at the Children’s Hospital of Philadelphia (*6*) (Supplementary Table 1). We first divided patients into two groups based on their initial response to the CAR T therapy: 10 patients (responders) had a complete remission defined by morphologic assessment of the bone marrow as M1 (<5% leukemic blasts), whereas the remaining 2 patients did not show an objective response to the therapy (non-responders, NR). We next subdivided responders into those who had a very durable complete remission (>54 months, CR, *n* = 5) and those who had a CD19-positive relapse during the course of the trial (RL, *n* = 5; median relapse-free remission duration = 9.6 months).

We sought to assess whether the intrinsic molecular characteristics of the infusion products could be predictive of clinical outcomes, reasoning that differences in T cell states and subtype proportions may determine both initial and long-term responses. Since the study of CAR T cells at baseline might not suffice to gain insights into the dynamics of early responses upon CAR engagement, we developed a stimulation method to emulate CAR-specific activation, for example, by B-ALL leukemic cells. To that end, we analyzed CAR T cells in infusion products upon CAR-specific stimulation with human CD19-expressing antigen presenting cells (APCs), and compared them to 3 additional conditions: TCR-mediated stimulation, non-target APC stimulation, and unstimulated controls (Fig. 1a). We engineered a NIH3T3 mouse cell line to stably express the human proteins CD19, CD86, and 4-1BB ligand (CD19-3T3 cells) (Supplementary Fig. 1a), and a control cell line expressing human mesothelin instead of CD19 (MSLN-3T3 cells) (Supplementary Fig. 1b). We co-cultured the infusion product of each patient with the engineered cell lines for 6 hours at a 1:1 ratio. In addition to these experiments, we stimulated the same infusion batches using anti-CD3/CD28-coated beads for the same duration to examine the TCR-mediated activation states.

**Figure 1.**
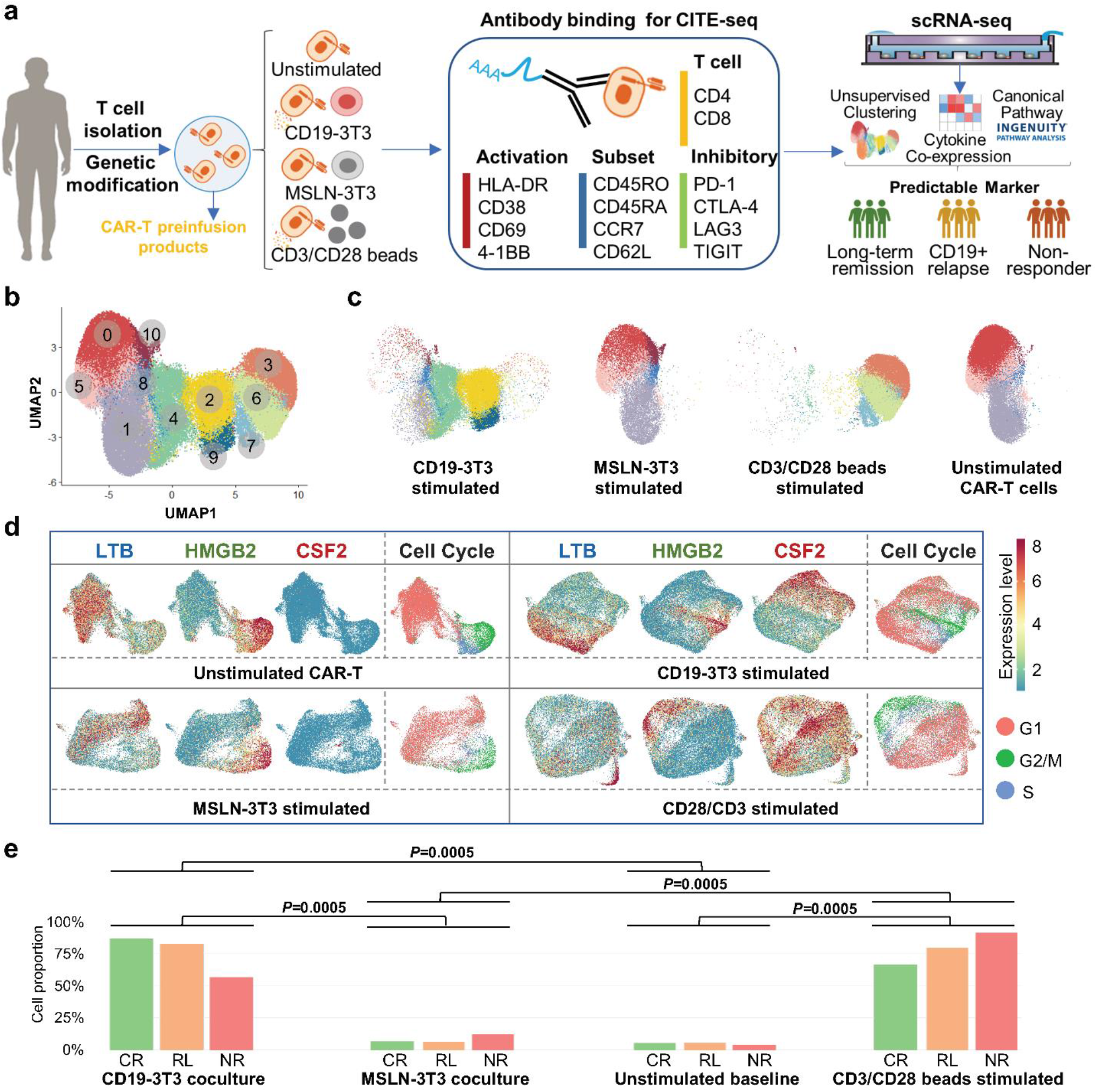
A functional landscape and activation states of CD19 CAR-T pre-infusion products from patients with ALL. (**a**) Schematic of experimental design. MSLN, mesothelin. (**b**) UMAP plot of 97,981 single CAR-T cells collected from 12 patients. 11 clusters are identified through unsupervised clustering. (**c**) UMAP distribution of all profiled cells separated by stimulation conditions. (**d**) The distribution of LTB, HMBG2, CSF2 and cell cycle expression pattern at four experimental conditions. Color bar represents normalized expression level. (**e**) Quantification of fully active CSF2(+) cell proportion in three response groups at the four conditions. The *P* values were calculated with Wilcoxon signed-rank test.

We performed scRNA-seq of the CAR T infusion products of the 12 patients under these 4 experimental conditions using a microwell-based approach (*29*). We adapted CITE-seq to this platform to concurrently profile 14 surface proteins through sequencing of antibody-derived DNA tags (ADT), including T cell markers (CD4, CD8), differentiation and subtype markers (CD45RO, CD45RA, CCR7, CD62L), activation markers (HLA-DR, CD69, CD38, 4-1BB), and co-inhibitory markers (PD-1, CTLA-4, LAG-3, TIGIT). In our co-culture experiments, CD19-3T3 cells formed strong immune synapses with CAR T cells to induce CAR-specific activation. To separate the transcriptome of CAR T cells and APCs, we mapped the sequencing reads to the concatenation of the CAR construct, human, and mouse reference genomes, and classified each read based on the genome it best aligned to.

### A global transcriptomic landscape of single CAR T cell activation states

In total, we profiled the transcriptome of 101,326 cells from CAR T infusion products, out of which 97,981 cells were analyzed after quality control and filtering. Integrating and clustering the single-cell RNA-seq data across all patients and conditions identified 11 subpopulations with distinct transcriptomic signatures, which we visualized using UMAP (Fig. 1b). As expected, stimulated cells separated from their unstimulated or control counterparts in this representation and MSLN-3T3 co-cultured cells overlapped with unstimulated cells (Fig. 1c), indicating experimental consistency and a minimal impact of antigen-negative costimulatory-ligand-expressing mouse APCs. We performed Ingenuity Pathway Analysis (IPA) (*30*) to explore the biological themes associated with the differentially expressed genes (DEGs) of each cluster, and found CAR-T cells in these subpopulations were highly heterogeneous in terms of their basic cell programs, cell cycle regulations, metabolic activities, T cell activation spectrums and effector functions (Supplementary Fig. 2).

Differential expression analysis between each subpopulation and all other cells revealed that subpopulations associated with stimulated cells (clusters 2, 3, 4, 6, 7, and 9) had significant (q-value < 0.05) upregulation of *CSF2* (colony stimulating factor 2, GM-CSF), *IFNG* (interferon γ, IFN-γ), and *GZMB* (granzyme B), and downregulation of *CCL5* (C-C motif chemokine 5), *CD52, LTB* (lymphotoxin β), and *STMN1*, suggesting that CAR T cells in these subpopulations are active functional effectors (Supplementary Fig. 3a). Two C-C motif chemokines, *CCL3* (MIP-1α) and *CCL4* (MIP-1β), and two C motif chemokines, *XCL1* and *XCL2*, were also upregulated in 5 of these clusters. In addition, cells co-cultured with CD19-3T3 APCs separated from anti-CD3/CD28-stimulated cells in the UMAP representation (Fig. 1c), suggesting differences in the gene expression programs induced by TCR and CAR-mediated T cell activation. Consistent with this observation, we found a deficit of *IL13* and *IL2* expression in subpopulations enriched for TCR-stimulated cells (clusters 3, 6 and 7), among other differences (Supplementary Fig. 3b).

### The CAR T cell activation states and their association with initial treatment responses

The presence of lentiviral vector elements in the CAR construct allowed us to differentiate the expression of the CAR from that of homologous endogenous TCR and co-stimulatory elements (Supplementary Fig. 4a). Based upon the fraction of cells with detected CAR construct expression among active cells in the stimulated samples, we estimated that the sensitivity to detect the expression of the CAR construct in the scRNA-seq data was 47%. Using this estimate, we inferred that 30% (90% confidence interval (CI): 21-41%) of the T cells in the infusion products were CAR+ (Supplementary Fig. 4b), out of which respectively 57-68% and 34-53% (90% CI) were CD4+ and CD8+ (Supplementary Fig. 4c), in agreement with previous reports of CAR transduction efficiency(*6, 31*).

Based on these results, we restricted our analysis to only CAR+ cells in the infusion products and examined each experimental condition separately. Clustering and differential expression analysis of cells at baseline revealed two major populations of CAR T cells, characterized respectively by high expression levels of *HMGB2* (high mobility box group 2) and *LTB* (Fig. 1d and Supplementary Table 2). The population of *HMGB2*-expressing CAR T cells was marked by high expression of cell cycle genes (Supplementary Table 2), consistent with these cells being actively proliferating. In contrast, the population of *LTB*-expressing cells consisted mostly of cells in the G1 phase of cell cycle (Fig. 1d) and had expression of *CCL5* (Supplementary Table 2). The same analysis in the single-cell data from CAR or TCR stimulated conditions revealed the presence of the same *HMGB2+* and *LTB+* cell populations across all experimental conditions, although their size was smaller upon CAR or TCR stimulation. Additionally, in the experiments involving CAR or TCR stimulation, our analysis revealed a large population of cells with high expression of *CSF2* (Fig. 1d) and other cytokines, including *IFNG, IL2, IL13, CCL3* and *CCL4*, and *XCL1* and *XCL2* (Supplementary Table 2). Thus, upon stimulation with CD19-3T3 cells or anti-CD3/CD28 beads, the proportion of *CSF2+* CAR T cells significantly increased for all patients (Fig. 1e, mean fold change (MFC)=22.2 and 20.7 for respectively for CAR and TCR stimulation). This increase was not observed upon co-culture with MSLN-3T3 control cells, consistent with *CSF2* expression being induced by the CAR or TCR activation. Based on these results, we identified *CSF2+* CAR T cells as CAR T cells in a functionally active state.

Reasoning that differences in the number of functionally active CAR T cells upon stimulation might be related to disparities in the clinical responses, we compared the proportion of *CSF2*+ cells between the clinical response groups defined above. However, we did not find any significant difference in the abundance of *CSF2*+ cells between the 3 groups of patients (Fig. 1e), although we did observe a significant anti-correlation between the duration of the response to the therapy and the proportion of *HMGB2+* CAR T cells upon stimulation with 3T3-CD19 cells (Spearman’s *r* = −0.65, *p*-value = 0.02).

### Failure to sustain a long-term response to the therapy is associated with a deficit of Th2 function in antigen-specific CAR T activation

We next investigated the heterogeneity of activation states of functionally active CAR T cells upon stimulation with CD19-3T3 cells, reasoning that differences among activation states might contribute to clinical responses. We performed an unsupervised analysis to identify modules of co-expressed cytokines (Methods). Our analysis of CAR-stimulated cells identified multiple functional cytokine co-expression modules, including two type-1-like cytokine modules comprising an array of type-1 response genes including *IFNG* and *IL2*, a type-2 cytokine module consisting of *IL4, IL5, IL9, IL13*, and *IL31*, and a module consisting of beta chemokines like *CCL3, CCL4* and related factors (Fig. 2a). A module consisting of *LTB, CCL5, IL16, IL32* and other cytokine genes related to the *LTB*-expressing cell subpopulation described above was also identified. Additionally, we considered a literature-derived expression module for terminally differentiated CD8 T cells (*32*), which was characterized by the expression of *ZEB2* transcriptional repressor. After removing cells that did not belong to these modules, we generated UMAP representations to delineate the distribution of each functional module, in which cells were located according to their inferred cytokine module activities (Fig. 2b). Expectedly, MSLN co-cultured control cells separated in the UMAP representation and showed high expression of the *LTB* cytokine module (Fig. 2c).

**Figure 2.**
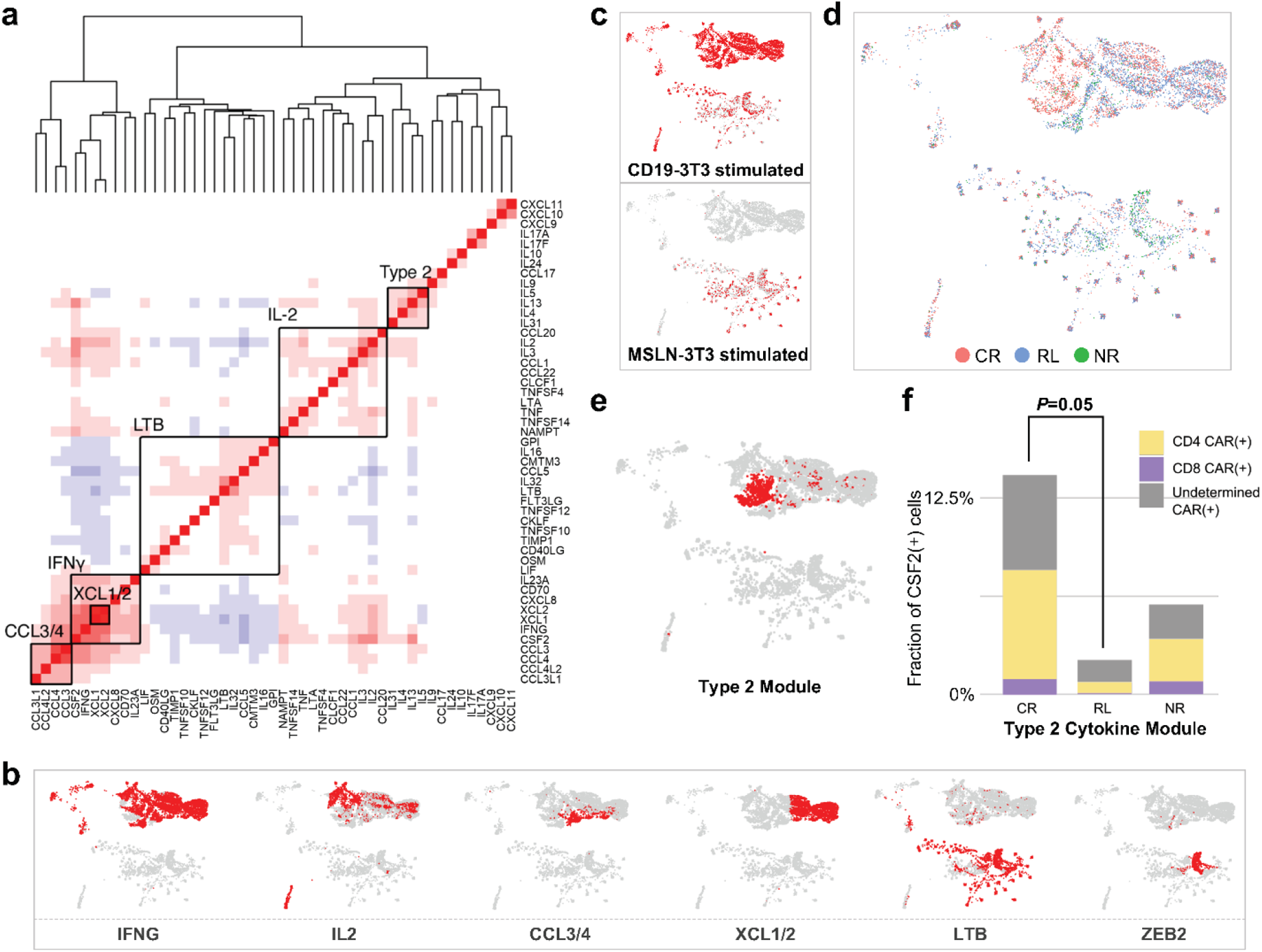
Co-expressed cytokine module analysis of CD19-specific stimulated CAR T cells identified functional heterogeneities and a deficit of Th2 function in CD19-positive relapsed patients. **(a)** Co-expressed cytokine modules identified in CD19-specific stimulated CAR(+) cells. Genes are ordered by hierarchical clustering. **(b)** The distribution of the identified modules on the UMAP. **(c)** An integrated cytokine module representation UMAP split by experimental conditions. **(d)** The localization of cells from each response group on the integrated UMAP. **(e)** The distribution of Th2 module. **(f)** Comparison of Th2 module-expressed CSF2(+) CAR(+) active cell proportion between very durable remission patients (CR) and CD19-positive relapsed patients (RL). The *P* values were calculated with Wilcoxon signed-rank test.

In the landscape of major responsive states, CAR+ cells from CR, RL, and NR patients were localized in the representation (Fig. 2d), suggesting a distinct cytokine module expression profile for these cells. Notably, the type-2 (Th2) module was found to be enriched in a region containing mostly cells from CR patients (Fig. 2e), implying that Th2 function might be required for maintaining a long-term remission in CAR T therapy. A quantitative comparison between the landscape of CAR T cells from NR, RL, and CR infusion products upon CAR-mediated stimulation identified a significant depletion of CAR+ cells expressing the type-2 cytokine module in RL patients compared to CR patients (MFC=4.14, Wilcoxon rank sum test *p*-value=0.05) (Fig. 2f). These data promoted more detailed investigations of the differences in the activation states between CAR T cells from CR and RL patients.

### Functional immune programs in activated CAR+ cells do not distinguish CD19-positive relapse from CR subjects, except for Th2 related pathways, genes and upstream regulators

To identify the characteristics of functional CAR T states associated with durable responses, we performed a global differential expression analysis between the CAR+ cells from CR and RL patients at unstimulated basal level or upon CAR-stimulation. The analysis of baseline CAR T cells only revealed the upregulation of EIF2 signaling in the CR group (Supplementary Fig. 5a,b), which might represent active protein synthesis and translation initiation in these cells (*33*). Comparing CAR T cells between CR and RL patients upon antigen-specific stimulation, showed the upregulation of genes associated with Th2 helper related cytokine production, such as *IL4, IL5* and *IL13*, in CR patients (Fig. 3a), consistently with our previous cytokine expression module analysis. Other immune pathways like Th1 cytokine production, T helper differentiation, and ICOS-ICOSL signaling were comparable between the two response groups (Fig. 3b). In line with these findings, a network analysis of canonical pathways, upstream regulators, and biological functions regulated by the differentially expressed genes inferred that CD28, ICOS, IL-3 and IL-25 were collectively operative in promoting the activation of the Th2 pathway in CAR T cells from CR patients (Fig. 3c). To explore the cascade of upstream regulators and assist the interpretation of the observed gene expression differences, we also performed IPA upstream analysis. The transcription regulator *FOXP3* and the cytokines *IL27, IFNG* and *IL10* were particularly active in cells from RL patients. By contrast, *GATA3*, a master transcription factor mediating the activation of Th2 signature genes, and several genes involved in T cell activation, including *NFKB1, NFATC2, TNF, CD28* and *CD69*, were upregulated in CR patients (Fig. 3d). To ensure that the above observations were not biased by any single subject, we assessed the expression of *IL13, IL5, IL4* and *GATA3*. The results of this analysis showed ubiquity of expression enrichment of the four genes in CR patients compared with relapsed subjects (Fig. 3e,f).

**Figure 3.**
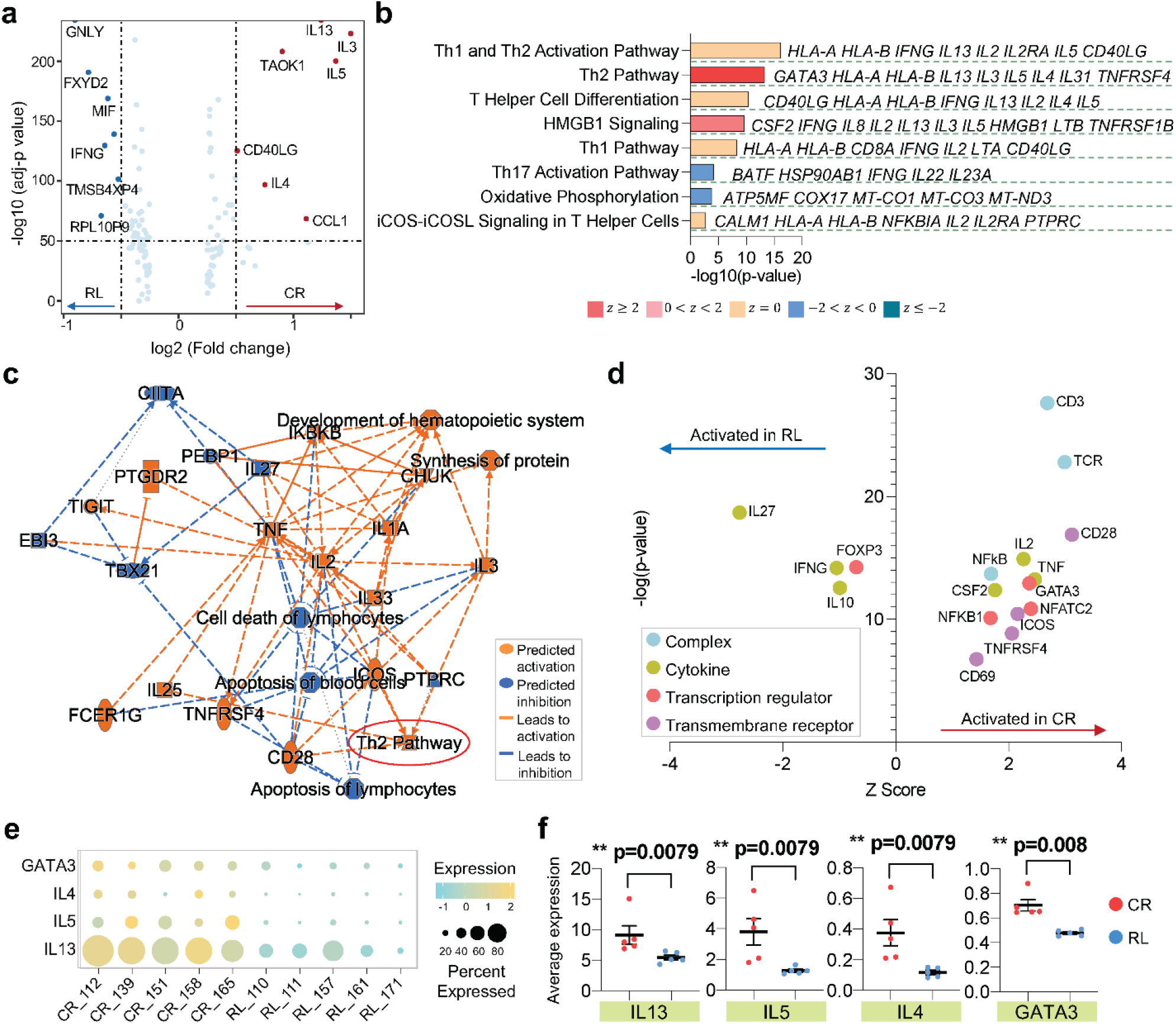
Th2 function related pathways, genes and upstream regulators are collectively downregulated in activated CAR T cells from patients with CD19-positive relapse. (**a**) Volcano plot of differentially expressed genes (DEGs) between CAR(+) cells from CR and RL patients. (**b**) Corresponding canonical pathways regulated by the highly differential genes identified in (**a**). Pathway terms are ranked by –log 10 (*P* value). The side listed gene names represent symbolic molecular marker related to the pathway. A statistical quantity, called *z* score, is computed and used to characterize the activation level. *z* score reflects the predicted activation level (*z* < 0, inhibited; *z* > 0, activated; *z* ≥ 2 or *z* ≤ −2 can be considered significant). (**c**) The graphical network of canonical pathways, upstream regulators, and biological functions regulated by DEGs identified in (**a**). (**d**) The predicted activation of upstream regulators, including complex, cytokine, transcription regulator and transmembrane receptor, in CR or RL patients. (**e**) Dotplot of Th2-related gene expression of each patient in CR and RL group. The size of circle represents proportion of single cells expressing the gene, and the color shade indicates normalized expression level. (**f**) The average expression level of genes *IL13, IL5, IL4*, and *GATA3* across all single cells in each patient and their comparison between CR and RL group. Each scatter point represents the average expression value of all single cells of specific patient. The *P* values were calculated with two-tailed Mann-Whitney test. Scatter plots show mean±s.e.m.

We next asked whether the proportion of cells with specific functional signature affect the sustainability of CAR T therapy. Clustering activated CAR+ cells from CR and RL samples identified 8 transcriptionally distinct populations (Fig. 4a). The top 3 genes differentiating each cluster indicated that CAR T cells in cluster 4 mainly functioned as Th2 helpers (Fig. 4b and Supplementary Fig. 6), and cells in clusters 2 and 3 had vigorous cytokine and chemokine gene expression including *IL2/3, IFNG, CSF2, XCL1/2* and *CCL3/4* (Fig. 4d). The cell proportion of cluster 4 was significantly elevated in CR patients and no fraction difference was observed for the combination of clusters 2+3 (Fig. 4c), suggesting that the lack of Th2 function rather than Th1 response could induce CD19-positive relapse. Consistently, the canonical pathway comparison between clusters did not detect differences in other major immune programs (Fig. 4e). To further check the CAR+ cell functional profiles at transcriptional level, we provided a single cell expression layout of all key immunologically relevant molecules across major categories, including effector, regulatory, stimulatory, inflammatory, and chemo-attractive functions (Fig. 4f). No apparent differences were found when comparing CR individuals with RL, with the exception of *IL4* and *IL13*. Together, these results show that, upon antigen-specific stimulation in CAR T infusion products, the Th2 pathway and upstream regulators are downregulated in RL patients as compared with their long-term CR counterparts, whereas other major immune programs remain comparable.

**Figure 4.**
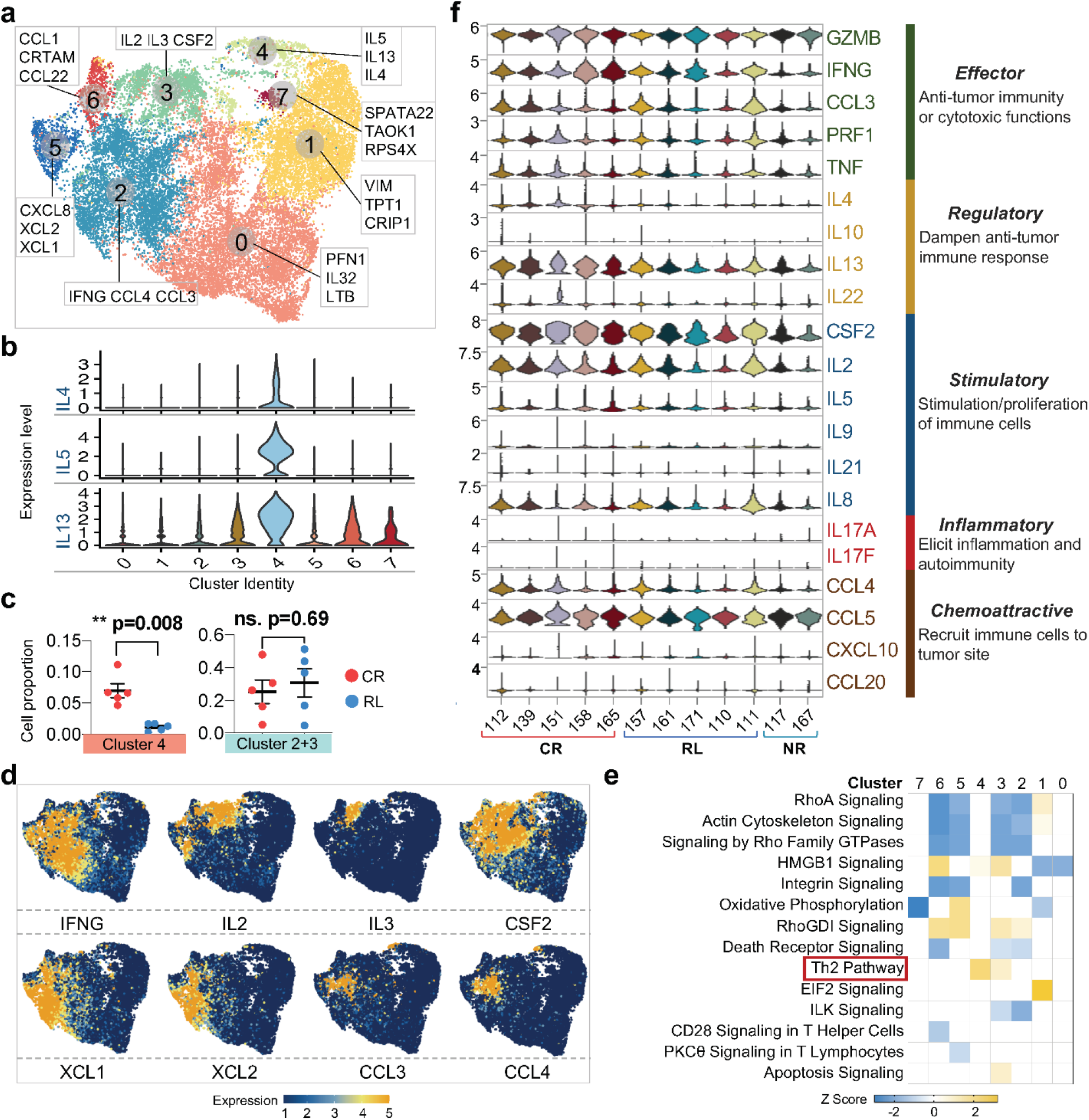
Single cell transcriptomic clustering of CAR T cells identifies comparable functional immune profiles between CR and RL patients, except for Th2 function. (**a**) UMAP re-clustering of CAR(+) cells from CR and RL patients and the top 3 dominant genes defining each cluster. (**b**) The expression of *IL4, IL5, IL13* in each identified cluster. (**c**) Comparison of cell proportion in particular cluster between CR and RL patients. The *P* values were calculated with two-tailed Mann-Whitney test. Scatter plots show mean±s.e.m. (**d**) The distribution feature plot of functional genes on the UMAP. (**e**) Canonical signaling pathway comparison between all the identified clusters. Differentially expressed genes of each cluster are used to identify the biological pathways. A statistical quantity, called *z* score, is computed and used to characterize the activation level. *z* score reflects the predicted activation level (*z* < 0, inhibited; *z* > 0, activated; *z* ≥ 2 or *z* ≤ −2 can be considered significant). (**f**) Single cell expression level violin plot of all key immunologically relevant cytokine genes from CAR(+) cells in each patient.

### Phenotypic proteomic profiling reveals that T_SCM_ and T_CM_ proportions, rather than activation spectra or co-inhibitory signatures, could predict CD19-positive relapse

Given the role of cellular stemness and memory formation of CAR T cells in mediating the efficacy and persistence of the therapy, we investigated the composition of naïve (T_N_), stem cell-like memory (T_SCM_), central memory (T_CM_), effector memory (T_EM_), and effector (T_EF_) subsets in the infusion products from the 3 clinical response groups, based on CD62L, CCR7, CD45RA and CD45RO cellular protein expression from our CITE-seq data (Supplementary Fig. 7) (*34*). For unstimulated baseline CAR T cells, we found significantly higher proportions of T_SCM_ and T_CM_ in the 5 CR patients, and a lower proportion of T_EM_ cells in these durable responsive cells (Fig. 5a,b). We also observed that a majority of CAR T cells derived from NR patients developed late memory signatures (Fig. 5a). Upon CAR stimulation, the lower abundance of T_SCM_ and T_CM_ cells was still significantly associated with the relapsed subjects (Fig. 5c,d). There was no difference in the frequency of T_EM_ CAR T cells, yet a higher ratio of cells in the RL group differentiated into a T_EF_ state compared with the CR group. Additionally, the early memory-to-effector (T_SCM_+T_CM_ to T_EF_) ratio was the lowest in NR patients among the 3 response groups (Fig. 5c). The expression of memory-related molecule CCR7 explained the disparities of CAR T differentiation subsets composition between long-term responders, responsive patients with relapse, and non-responders, both at baseline and upon CAR stimulation (Fig. 5e).

**Figure 5.**
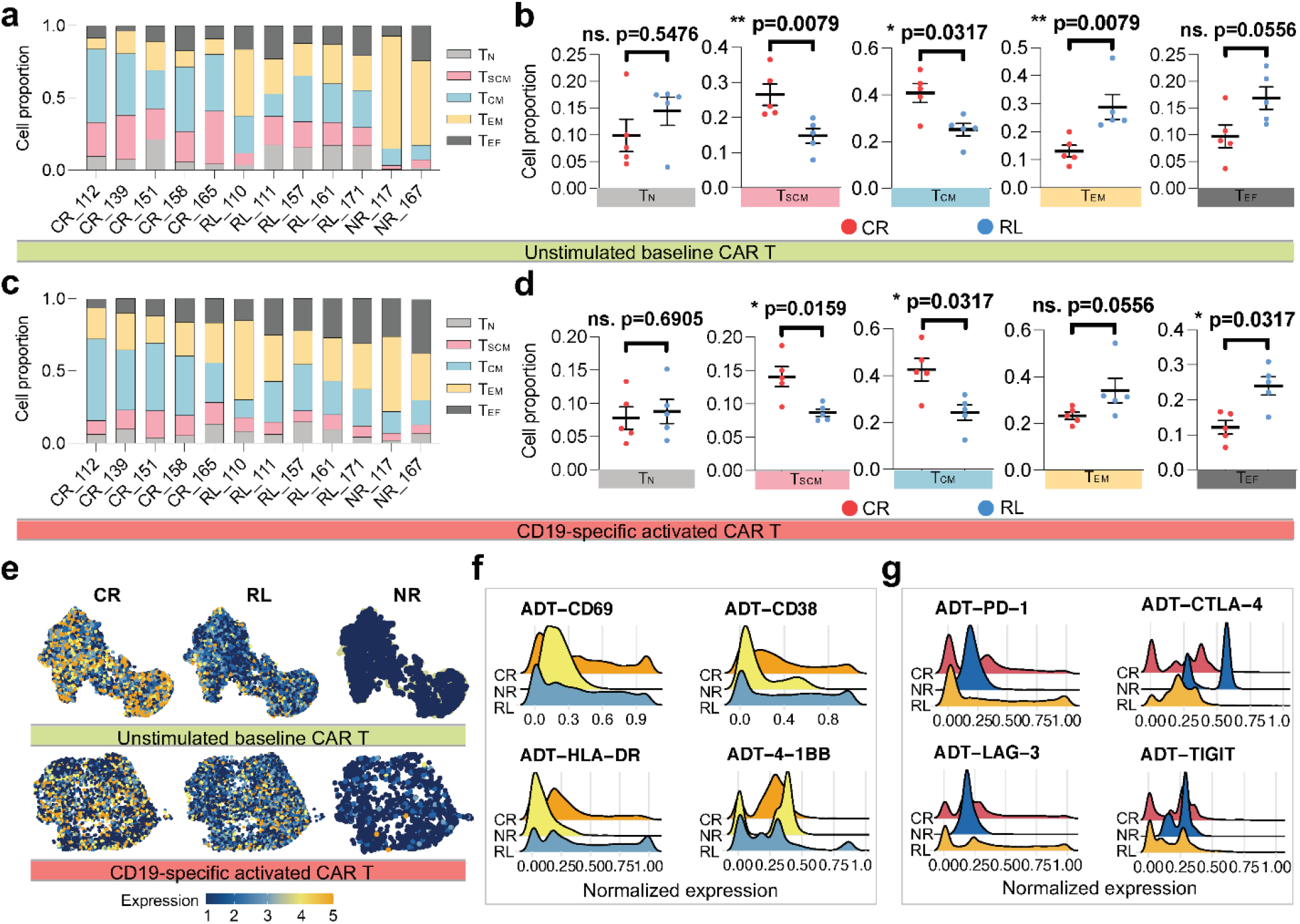
Phenotypic proteomic profile reveals the dampened capacity of maintaining early memory T cell states in CAR T cells from relapsed patients. (**a**) Differential state composition of unstimulated baseline CAR T cells in each patient, based on the expression of CD62L, CCR7, CD45RA and CD45RO from CITE-seq data. T_N_, naïve; T_SCM_, stem cell-like memory; T_CM_, central memory; T_EM_, effector memory; T_EF_, effector T cells. *x* axis represents responsive state and patient ID. (**b**) Proportion comparison of unstimulated CAR T cell phenotypic subsets between CR and RL group. (**c**) Differential state composition of CD19-specific activated CAR(+) cells in each patient. (**d**) Proportion comparison of activated CAR(+) cell phenotypic subsets between CR and RL group. (**e**) Feature plot of cellular protein CCR7 in unstimulated or activated CAR T cells, split by responsive groups. (**f**) Single cell expression distribution of the T cell activation related surface protein markers of each group. (**g**) Single cell expression distribution of co-inhibitory surface protein markers of each group. The *P* values were calculated with two-tailed Mann-Whitney test. ns. Not significant. Scatter plots show mean±s.e.m.

We also evaluated the surface protein markers of T cell activation and co-inhibition in activated CAR+ cells across patient groups. Of the 4 major activation markers, a higher proportion of cells from NR patient had a relatively low expression of CD69, CD38 and HLA-DR, whereas the expression levels were not significantly different between the CR and RL groups (Fig. 5f and Supplementary Fig. 8a,b). Likewise, the genes encoding these activation markers were expressed at similar levels (Supplementary Fig. 8c,d). In the evaluation of T cell co-inhibitory signatures associated with dampened proliferation and effector functions, namely PD-1, CTLA-4, LAG-3, TIGIT and their corresponding genes, we did not observed differences between CR and RL patients (Fig. 5g and Supplementary Fig. 9a-d). In summary, this phenotypic analysis indicates that CD19-positive relapse is associated with deficiencies in the capacity of CAR T cells for maintaining T_SCM_ and T_CM_ states.

### CAR T cells from NR patients fail to mount robust type 1 responses and have a higher proportion of terminally differentiated CD8 CAR T cells

The analysis of cytokine module expression described above revealed some commonalities between the two NR patients in our cohort which, although not statistically significant, might represent more general cellular characteristics associated with the lack of initial response to the therapy. In particular, infusion products of NR patients upon CAR stimulation showed a depletion of CD4+ and CD8+ CAR T cells expressing type-1 cytokine modules as compared to other patients (Fig. 6a, Wilcoxon rank sum *p*-value=0.06, 0.06, 0.12 and 0.12, respectively for the *XCL1/2, CCL3/4, IL2* and *IFNG* cytokine modules). In contrast, cells from NR patients were enriched for terminally differentiated CD8 cells expressing the *ZEB2* module (Fig. 6a, MFC=8.3, Wilcoxon rank sum test *p*-value=0.03). These results suggest that the lack of robust initial response to the therapy might be associated with a failure of stimulated CAR T cells to mount a robust type 1 adaptive response and an enrichment for terminally differentiated CD8 CAR T cells primed toward a low-cytokine expression state. Consistent with this hypothesis, we observed a significant correlation between the duration of the clinical response to the CAR T cell therapy and the proportion of CAR T cells expressing type-1 cytokine modules (Spearman’s *r*=0.76 and 0.65, *p*-value=0.004 and 0.02, respectively for the *CCL3/4* and *IL2* modules), and an anti-correlation between the proportion of CAR T cells expressing the *ZEB2* cytokine module (Spearman’s *r*=-0.67, *p*-value=0.02), upon *ex vivo* CAR stimulation (Fig. 6b).

**Figure 6.**
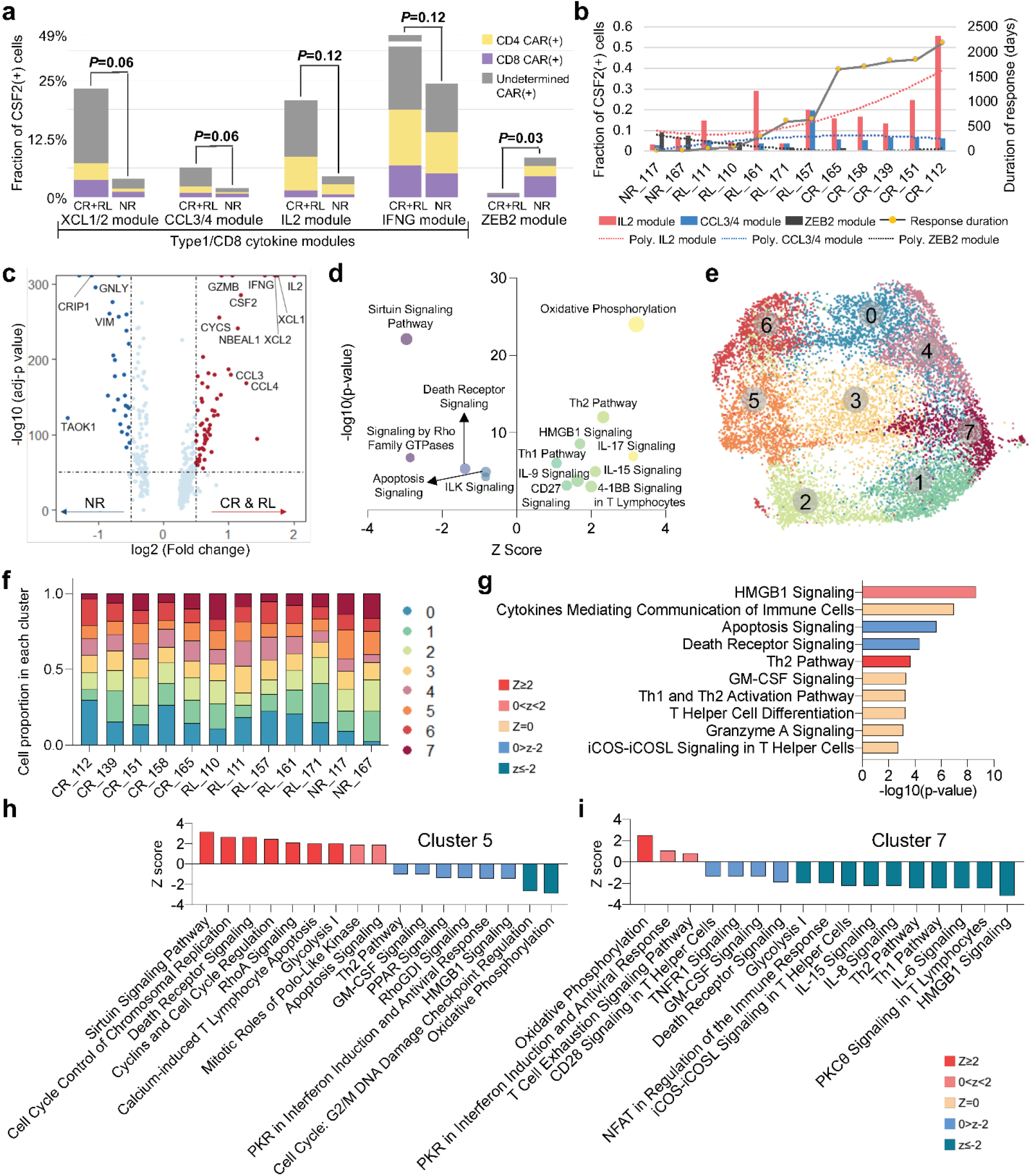
Co-expressed cytokine module and pathway analyses reveal attenuated effector functions of CAR T cells from non-responders. (**a**) Quantification of type-1 and *ZEB2* module expression proportions in active CSF2(+) CAR(+) cells between NR non-responders and other patients. The *P* values were calculated with Wilcoxon signed-rank test. (**b**) The correlation between *IL2, CCL3/4* and *ZEB2* module-expressed cell proportion and the duration of the clinical response to the therapy. The second-order polynomial fitting curves were computed to show the proportion changing trend. (**c**) Volcano plot of differentially expressed genes between responsive CR/RL patients and NR patients. (**d**) Corresponding canonical pathways regulated by the highly differential genes identified in (**c**). (**e**) UMAP clustering to identify transcriptional distinct clusters of CD19-specific activated CAR(+) cells. (**f**) Fraction of cells from each patient sample in each cluster. *x* axis indicates responsive state and patient ID. (**g**) Representative pathways identified in cluster 0. (**h**,**i**) Activated or inhibited pathways identified in cluster 5 (**h**) and 7 (**i**). *z* score reflects the predicted activation level (*z* < 0, inhibited; *z* > 0, activated; *z* ≥ 2 or *z* ≤ −2 can be considered significant).

To characterize the gene expression profile of CAR T cells from NR patients upon CAR-mediated stimulation, we performed differential gene expression analysis between the CAR+ cells from NR patients and all the other patients in the cohort (Fig. 6c). This analysis revealed that cytokine genes such as *IL2, CSF2, IFNG, XCL1/2* and *CCL3/4* were downregulated in NR patients, in agreement with the findings from our previous cytokine module analysis. IPA of these differentially expressed genes confirmed a strong activation of HMGB1, Th1, Th2, IL-9, IL-15, IL-17, 4-1BB and CD27 signaling in responders as compared to NR subjects, whereas apoptosis and death receptor signaling were upregulated in cells from NR patients (Fig. 6d). Clustering CAR+ cells identified 8 subpopulations with distinct gene signatures (Fig. 6e and Supplementary Fig. 10). We observed NR subjects had the lowest proportion of cells in cluster 0 (Fig. 6f), in which T cell functional pathways such as HMGB1 and Th2 signaling were highly activated (Fig. 6g). The majority of T cell activation and effector pathways were inhibited in clusters 5 and 7, whereas calcium-induced T lymphocyte apoptosis, glycolysis and T cell exhaustion signaling were active (Fig. 6h,i). As expected, cells of NR patients were enriched in these two clusters (Fig. 6f). Taken together, these results suggest that a deficit of CAR T cell activation to elicit multiple immune effector functions is associated with poor initial anti-leukemic responses in NR patients.

### TCR-mediated bead stimulation of infusion products fails to capture clinical responses

The findings described in previous sections show that the landscape of CART19 activation states upon CAR-mediated *ex vivo* stimulation is associated with the clinical response to the therapy. To assess whether stimulation via anti-CD3/CD28-beads was sufficient to uncover the determinants of response, we used the same computational approach to search for differences in the gene expression programs associated with the CAR T cells of CR, RL, and NR patients upon TCR-mediated stimulation (Fig. 7a,b). This analysis revealed that, overall, the cytokine co-expression modules were consistent with those identified in CD19-stimulated CAR T cells, except for a Th17-like module consisting of *IL17F, IL17A, IL22, IL26*, and *CCL20* that was uniquely observed upon TCR-mediated simulation (Fig. 7a). Nevertheless, the number of cells expressing this module was extremely low (Fig. 7c). We also observed a type-1 module in correlation with the expression of the apoptosis antigen ligand *FASLG*, possibly reflecting substantial activation-induced cell death upon bead stimulation. However, contrary to what we observed in the case of CAR-mediated stimulation, we did not find any significant difference between CR, RL, and NR patients in the proportion of cells expressing each cytokine co-expression module upon TCR-mediated stimulation (Fig. 7d), suggesting that this stimulation mechanism was incapable of uncovering the biology of CAR T cells that underlies differences in clinical responses. Consistently, we did not observe any difference on the gene expression levels of major cytokines across the three response groups (Supplementary Fig. 11a,b). Similarly, IPA showed an overall consistency between the main pathways identified in the two conditions, although the activation/inhibition level and cell number of specific pathways differed substantially (Supplementary Fig. 12a-d).

**Figure 7.**
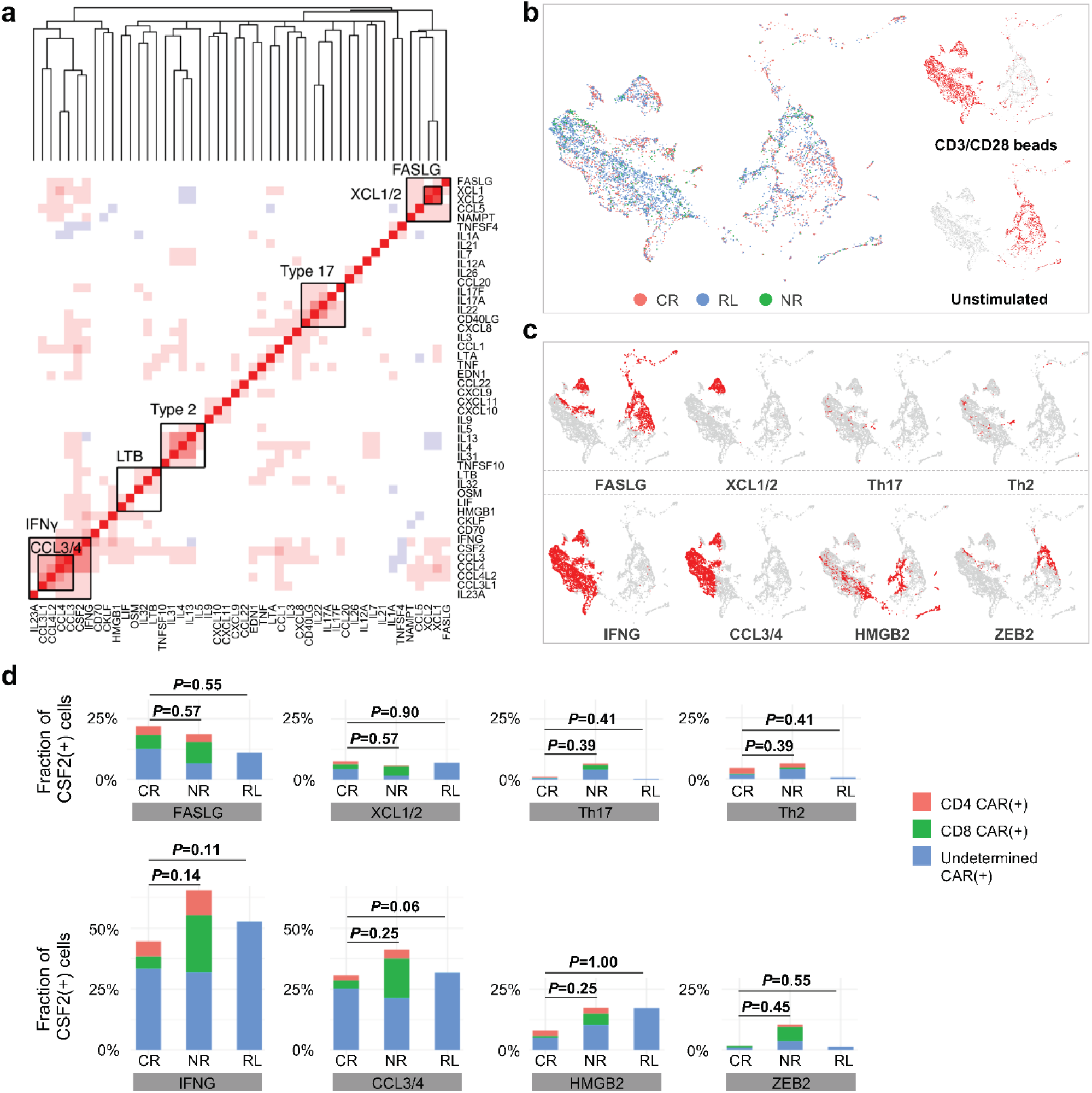
TCR-mediated anti-CD3/CD28-beads stimulation of CAR T cells fails to predict clinical responses. (**a**) Co-expressed cytokine modules identified in TCR-mediated stimulation. Genes are ordered by hierarchical clustering. (**b**) An integrated cytokine module representation UMAP. Cells are colored by responsive states and UMAPs are split by experimental conditions. (**c**) The distribution of the identified modules on the UMAP. (**d**) Quantification of identified modules between CR, NR and RL patients. The *P* values were calculated with Wilcoxon signed-rank test.

## Discussion

CART19 cell therapy is highly effective in pediatric and adult patients with B-ALL with 90% patients responding initially to the therapy (*6*). Despite this high response rate, relapse occurs in nearly 50% of patients, among which a large fraction is CD19 positive (*8-10*). Owing to the costly and lengthy process to manufacture and administer CAR T therapy in a patient-specific fashion, predictive biomarkers are urgently needed to better weigh the cost and value of treatment and specific strategies are expected to ameliorate CAR T products to effectively extend the remission durability. Significant efforts have been made toward identifying patient characteristics, including tumor burden and immune status, in order to correlate with efficacy and/or toxicities of CAR T therapy (*7, 35*). Early expansion of peripheral blood CAR T-cells *in vivo* has been associated with favorable therapeutic outcome (*7, 11*), but it does not able to predict the response or relapse prior to infusion. Furthermore, to date, the mechanisms inducing CD19-positive relapse remain an open question. Since the dose of CAR T cells approved as a standard of care for B-cell malignancies can vary up to 10-fold and the sustained remission is largely independent of dosage in long-term follow up studies (*6*), the composition of the infusion product appears to be more critical than the quantity of CAR-transduced T cells. Thus, it is crucial to understand if the characteristics of infused CAR T cells contribute to the CD19-positive relapse and therapeutic failure.

We conducted single-cell transcriptional profiling of CD19-targeting 4-1BB/CD3-signaling CAR T cells in infusion products of B-ALL patients upon stimulation with human CD19-expressing APCs, and observed significant intrinsic functional heterogeneity. Through unbiased analyses of distinct co-expressed cytokine modules on a fully activated CAR T subset marked by high *CSF2* expression, we identified two type-1-like cytokine modules comprising an array of Th1 response genes including *IFNG* and *IL2*, a type-2 cytokine module consisting of *IL4, IL5, IL9, IL13*, and a module consisting of beta chemokines like *CCL3, CCL4*. A partially activated subset marked by the expression of *LTB* and other cytokines like *CCL5, IL16*, and *IL32* was also found, which constituted the dominant feature for MSLN co-cultured CAR T-cells cells. Single-cell multiplexed cytokine profiling was previously utilized to characterize healthy donor 4-1BB/CD3 signaling CART19 cells stimulated with CD19-coated beads and showed highly heterogeneous functional states (*36*). The most poly-functional subset produces GM-CSF, the protein encoded by *CSF2*, type-1 cytokine IFN-γ, type-2 cytokine IL-13, and beta chemokines like MIP-1α and MIP-1β, encoded by *CCL3* and *CCL4* genes, respectively, in both CD4 and CD8 CAR T cells, which is in general agreement with what we observed in this study.

Comparing the co-expressed cytokine expression profile of CD19-positive relapsed patients with that of durable responders, we found that CAR T cells from RL patients had a significantly deficit of Th2 module expression. Differential gene expression analyses identified that all major genes encoding Th2 functions, including *IL13, IL5, IL4* and the upstream regulator *GATA3* were upregulated in long-term CR patients compared with RL patients. In contrast, these analyses did not identify differences in the expression of Th1 or beta chemokine response related genes, suggesting the need for both type-1 and type-2 functions to achieve a sustained long-term remission. These findings suggest a potential strategy to improve the therapeutic outcome by re-engineering the infusion product with augmented type-2 signaling or by booting Th2 functions post infusion.

Recently, Deng *et al*. performed scRNA-seq of autologous axi-cel anti-CD19 CAR T cell infusion products from 24 patients with LBCL (*20*). Their data revealed three-fold higher frequencies of CD8 T cells expressing memory signatures in patients who achieved a complete response than patients with partial response or progressive disease. Moreover, CD8 T cell exhaustion was associated with a poor molecular response, suggesting both T cell memory status and immune function capacity in infusion products can contribute to therapeutic outcome. By utilizing our CITE-seq surface proteomic data, we identified T cell differentiation subsets and found that CAR T cells from relapsed patients had significantly lower proportion of T_SCM_ and T_CM_ early memory states under both unstimulated baseline and CD19-specific activated conditions. However, no apparent differences of activation- or co-inhibitory-related marker expression were observed between CR and RL group. Further prospective investigation in a larger cohort will be indispensable to fully validate this observation.

We also found that the inability for CAR T cells to mount a robust type 1 cytokine module response upon CAR-mediated stimulation was associated with non-responders. Terminally differentiated CD8 T cells characterized by the expression of *ZEB2* transcriptional repressor were significantly correlated with the failure of initial response (*32*), which may represent a potential target for combination therapy to improve the initial response rate. We also found that T lymphocyte apoptosis, glycolysis and T cell exhaustion signaling were active in a high fraction of CAR T cells from NR patients, and our phenotypic data showed that the early memory-to-effector ratio in activated NR CAR T cells was the lowest among the 3 response groups. These results are in agreement with the results of a previous study, where we conducted transcriptomic profiling of CAR T cells in patients with CLL and found memory-related genes, including IL-6/STAT3 signatures, were enriched in responders whereas T cells from non-responders upregulated programs involved in effector differentiation, glycolysis, exhaustion and apoptosis (*21*).

While our study provided a single-cell transcriptional landscape of CAR T cell activation states upon antigen specific stimulation and identified subsets with distinct gene signatures associated with therapeutic response, it is yet to be evaluated in CAR T cells that include other costimulatory domains or target antigens other than CD19 or diseases other than B-ALL. The spectrum of effector functions may vary as a function of the co-stimulatory domain in the CAR construct. Rossi et al. conducted single-cell cytokine analysis of CD19 CAR T cells in the infusion product engineered with the CD28 co-stimulatory structure and observed polyfunctional T cell subsets capable of deploying multiple immune programs including IFN-γ, IL-17A, IL-8, and MIP-1α correlate significantly with objective responses in patients with Non-Hodgkin’s Lymphoma (NHL), which was the first study to show single-cell immune function heterogeneity in infusion products has a major association with clinical outcomes of CAR T therapy (*37*). Notably, the polyfunctional T cells also required type 1 functions (IFN-γ and TNF-α) and beta chemokines (MIP-1α and MIP-1 β), supporting the hypothesis that transfer of a population of cells inclusive of multiple T-cell subsets could better support the production of CAR T cells capable of diverse, polyfunctional and immune activities, consistent with our observation in patient 4-1BB CAR T cells. However, the polyfunctional cells showed little type-2 functions except IL-5 in a less polyfunctional subset co-expressing TNF-α, in contrast to co-expressed type-1 and 2 modules in the fully activated subset in our study. In addition, polyfunctional IL-17A-producing T cells were identified in strong correlation with antitumor efficacy as well as neurotoxicity whereas IL-17 was undetectable at both transcriptional and protein levels in 4-1BB CAR T cells from patients or healthy donor, highlighting the existence of functional differences between 4-1BB and CD28 CAR T products. While we observed the long-term persistence and sustained remission correlates with type-2 functions (i.e., *IL13*) in ALL patient infusion products, it is unclear if it is also required for CD28 CAR T cells although they were found to be less persistent than 4-1BB CAR T cells (*17*).

Single-cell transcriptome profiling allowed for the dissection of CAR T cell activation states at the transcriptional level and identified the immune functional programs associated with both short term and long-term therapeutic efficacy. However, the underlying mechanisms are yet to be explored in highly heterogeneous CAR T infusion products. Specific to CAR T cell engineering, the variation of CAR integration sites in individual T cells may contribute to functional heterogeneity of CAR T cells and ultimately the outcomes of treatment. Nobles *et al*. investigated clonal population structure and therapeutic outcomes in CLL patients by high throughput sequencing of vector integration sites and found genes at integration sites enriched in responders were commonly found in cell-signaling and chromatin modification pathways, suggesting the insertional mutagenesis in these genes can be linked to treatment outcomes (*38*). Single cell level integration site profiling in conjunction with co-measurement of transcriptome, if developed, may further elucidate the impact of insertion site variation on CAR T cell heterogeneity. Recently, Wang *et al*. reported on a method for joint profiling of chromatin accessibility and integration sites of CAR T cells at population and single cell level, which may address a question whether CAR T integration at certain genomic regions can rewire the regulatory landscape of T cells (*39*), but it is yet to be applied to patient derived CAR T cells in correlation with clinical outcomes.

An outstanding question is if the functional heterogeneity and the activation modules of CAR T cells in the infusion product can be solely attributed to the characteristics of the patient immune cells to start with or can be improved through optimized manufacturing process by analyzing apheresis products in comparison with corresponding infusion products. We observed that neither unstimulated nor TCR-stimulated CAR T cell characteristics was associated with therapeutic outcomes, but only CAR specific signaling and cytokine responses elicited by antigen-specific stimulation could predict CR, RL and NR, indicating the necessity to characterize infusion products to develop more informative and accurate predictive biomarkers. In-depth analysis of proteomic (surface epitopes and cytokines), transcriptional (mRNAs), genomic (i.e., integration sites), and epigenomic (chromatin state) landscapes of single CAR T cells throughput the manufacturing process may further unveil the mechanisms of their function changes with time and inform new strategies to improve manufacturing or control the quality of autologously transferred CAR T cells for individual patients. Finally, the clonal expansion *in vivo* post infusion constantly alters the heterogeneity of circulating CAR T cells in patients (*40*) and a finite number of clones are the major effector cells inducing therapeutic remission and long-term surveillance to completely eliminate tumor cells. However, it remains to be revealed how the immune function landscape correlates with clonal evolution and the persistence in patients throughout the treatment and the long-term remission.

## Methods

### Patient samples

Samples were acquired from patents with relapsed/refractory B-ALL who enrolled in pilot clinical trials designed to assess the safety and feasibility of CD19-dericted chimeric antigen receptor (CTL019) T cell therapy, which was conducted at the Children’s Hospital of Philadelphia (ClinicalTrials.gov number, NCT01626495) and the University of Pennsylvania (ClinicalTrials.gov number, NCT01029366). Written informed consent for participation was obtained from patients or their guardians according to the Declaration of Helsinki. All laboratory operations were under principles of International Conference on Harmonization Guidelines for Good Clinical Practice with established Standard Operating Procedures (SOPs) and protocols for sample receipt, processing, freezing, and analysis. All ethical regulations were strictly followed.

### T cell isolation, vector production, and generation of CTL019 cells

Autologous peripheral blood mononuclear cells (PBMCs) were collected by standard leukapheresis. T cells were enriched by mononuclear cell elutriation, then washed and activated with anti-CD3/CD28-coated paramagnetic beads. A lentiviral vector containing a previously described CD19-specific CAR with 4-1BB/CD3ξ transgene was constructed and produced (*31*), which was then used to transduce the cells during activation and was washed out 3 days after the culture initiation (*41*). A rocking platform device (WAVE Bioreactor System) was used to expand cells for 8 to 12 days, and the beads were then magnetically removed. CTL019 cells were harvested and cryopreserved in infusible medium. The final product release criteria are listed as following (*6*): cell viability ⩾ 70%, CD3+ cells ⩾ 80%, residual paramagnetic anti-CD3/CD28-coated paramagnetic beads ⩽ 100 per 3×10^6^ cells, Endotoxin ⩽ 3.5 EU/mL, Mycoplasma negative, Bacterial and fungal cultures negative, residual bovine serum albumin ⩽ 1 μg/mL, VSV-G DNA as a surrogate marker for replication competent lentivirus ⩽ 50 copies per μg DNA, transduction efficiency by flow cytometry ⩾ 2%, transduction efficiency by vector DNA sequence 0.02 to 4 copies per cell.

### Generation of human CD19- and Mesothelin NIH3T3 cells

NIH3T3 mouse fibroblasts were originally obtained from the American Type Culture Collection (ATCC) and were cultured in Dulbelcco’s modified Eagle’s medium (DMEM; Gibco), with glutamate and supplemented with 10% fetal bovine serum (FBS; Gibco) in a humidified incubator (37°C in an atmosphere of 5% CO2). The cells were transduced with a lentiviral vector encoding human CD19 (3T3-CD19), and a negative control cells were NIH3T3 cells expressing mesothelin (3T3-MSLN). Cells were sorted on a FACSAria (BD) to reach a >99% purity after transgene introduction. Mycoplasma and authentication were routinely performed before and after molecular engineering.

### In vitro coculture assay

CTL019 cells were thawed and cultured in OpTmizer™ T-Cell Expansion Basal Medium supplemented with OpTmizer™ T-Cell Expansion Supplement and GlutaMAX™ Supplement (Gibco) in the humidified incubator on day 1 for overnight rest. On day 2, dead cells were removed using EasySep™ Dead Cell Removal Kit (Stemcell) as per the manufacturer’s instructions, and certain number of cells were counted using hemocytometer before coculture assays. For the stimulation with 3T3-CD19 cells or 3T3-MSLN cells (target cells), 1×10^6^ CTL019 cells were mixed with same number of target cells and resuspended in 2mL medium; for the stimulation with anti-CD3/CD28 beads, 1×10^6^ CTL019 cells were mixed with 3×10^6^ beads and resuspended in 1mL medium; for the evaluation of unstimulated condition, 1×10^6^ cells were prepared. All the suspensions were cultured in RPMI medium (Gibco) containing 10% FBS in a tissue culture-treated 48-well plate (Fisher Scientific) for 6 hours in the incubator. After coculture, the anti-CD3/CD28 beads were removed by passage over a magnetic field.

### Cell staining with DNA-barcoded antibodies for CITE-seq

After 6h coculture, for each condition, cells were resuspended in 100 µl Cell Staining Buffer (Biolegend) added with 5 µl of Human TruStain FcX™ Fc Blocking reagent (Biolegend). Cell suspensions were then incubated at 4°C for 10 mins, during which the antibody-pool was prepared using 1µg of each TotalSeq™ antibody (Biolegend). After 30 mins incubation at 4°C, cells were washed 2 times with 1 mL PBS (Life Technologies) and finally resuspended in PBS at 5×10^5^ cells/ml for the downstream loading. TotalSeq antibodies for human T cell characterization panels: anti-CD4 (clone SK3), anti-CD8 (clone SK1), anti-CD45RO (clone UCHL1), anti-CD45RA (clone HI100), anti-CCR7 (clone G043H7), anti-CD62L (clone DREG-56), anti-HLA-DR (clone L243), anti-CD38 (clone HB-7), anti-CD69 (clone FN50), anti-4-1BB (clone 4B4-1), anti-PD-1 (clone EH12.2H7), anti-CTLA-4 (clone BNI3), anti-LAG3 (clone 11C3C65), anti-TIGIT (clone A15153G).

### Device Preparation and Microfluidic Operation of scRNA-seq

The detailed materials and methods for preparing and operating the scRNA-seq device have been described in our previous report (*29*). Briefly, the device consisted of a microwell array layer and a microfluidic channel layer, both of which were made by casting PDMS over the SU8 master wafers followed by degassing and curing at 80°C for 6-8 hours. After curing, PDMS was peeled off, and the two layers were cut to proper sizes and then plasma-bonded to assemble onto a glass slide. Prior to cell loading, device was pressurized to remove air bubbles inside the microwells using a manually operated syringe with outlet closed, and then primed for 1h at room temperature with 1% Bovine Serum Albumin (BSA, Sigma) in PBS. A total of 100µl cell suspension with a density of 5×10^5^ cells/ml was pipetted on the inlet reservoir and withdrawn into the device. Next, barcoded beads, lysis buffer and fluorinated oil were loaded sequentially to seal microwells. After cell and beads loading, the device was placed in a petri dish and exposed to three cycles of freeze-and-thaw, each of which included freezing at -80°C freezer or dry ice for 5 min followed by thawing at room temperature for 5 min. To capture mRNA onto beads, the device was then incubated for 1h inside an aluminum foil covered wet chamber. After incubation, the beads were retrieved by 6X saline-sodium citrate (SSC) buffer flushing. Finally, collected beads were washed twice with 6X SSC buffer and then proceeded to the reverse transcription step.

### scRNA-seq Library Preparation and Sequencing

Library preparations were performed as described in the DropSeq protocol (version 3.1) (*42*). Briefly, the captured mRNA was reverse-transcribed using Maxima H Minus reverse transcriptase (Thermo Fisher) with a custom template switching oligo. Then, the beads coated with cDNA were treated using Exonuclease I (Exo I, NEB) for 1h at 37°C with rotation to chew away any unbound mRNA capture probes. The cDNA was then amplified using a 13-cycle PCR whole transcriptome amplification. At this step, 0.4 µl of “additive” primers were added to cDNA PCR to increase yield of Antibody Derived Tag (ADT, cDNA derived from the TotalSeq™ antibodies) products. After the amplification, ADT-derived cDNAs (180bp) and mRNA-derived cDNAs were separated using SPRIselect (Beckman Coulter) beads at 0.6 ratio. After a standard Nextera tagmentation, PCR reactions (Nextera XT, Illumina) and another round of 0.6X purification using AMPure XP beads (Beckman Coulter), the mRNA library was ready to be sequenced. For ADT-derived cDNAs, 2X SPRI purifications and then an 8-cycle PCR were performed to amplify ADT sequencing library, followed by another round of 1.6X SPRI purification per manufacturer’s protocol. The quality of both the mRNA and ADT libraries were assessed by high sensitivity Bioanalyzer test (Agilent Inc.). To obtain sufficient read coverage for both libraries, we pooled 10% of ADT and 90% of cDNA library into one lane. The libraries were sequenced on HiSeq4000 (Illumina) at medium depth (average of 20k to 40k reads/cell) with 3 samples pooled into one sequencing lane.

### Processing of sequencing data

Reads from the mRNA libraries were aligned to the concatenated human (hg38) and mouse (mm10) reference genomes and the sequence of the transduced CAR construct (*31*) using STAR 2.6.1a. A UMI count matrix was built using the Drop-seq core computational protocol (version 2.0.0) (*42*). As reads were aligned to the concatenated human and mouse reference genomes, we were able to classify each uniquely aligned UMI as originating from a human or mouse cell. CITE-seq-Count was used to count UMIs in the ADT libraries. We filtered out cells with less than a specified total number of mRNA UMIs. This number was determined in a sample-specific manner by examination of log scale barcode rank plots, where a steep drop-off was taken to indicate the boundary between empty wells and wells containing a cell. This fraction was between 1×10^2.8^ and 1×10^3.2^ for all samples. For co-culture experiments we filtered out mouse cells by removing cells with more than a specified number of mouse associated UMIs. Again, this number was determined in a sample-specific manner by examination of plots of mouse vs. human UMI counts. After removing wells which contained a mouse cell from the analysis, residual background expression of mouse genes was removed by setting counts to zero. The mRNA expression data was log normalized using Seurat (*43*), and the Seurat v3 anchor-based integration workflow was used to integrate data from different experiments. We used 20 CCA dimensions and the default number of anchor neighbors and reduced the integrated mRNA data to 2 dimensions using UMAP for visualization (*44*). The ADT libraries were normalized using Seurat v3.

### Estimation of CAR+ cell fraction

To estimate the sensitivity to detect expression of the CAR, we counted the fraction of CAR mRNA expressing cells. We focused on cells from the CD19 APC co-culture experiments expressing CSF2, on the assumption that all CSF2 expressing cells had been successfully activated via CAR binding to CD19.To estimate the CAR+ fraction in the infusion samples we used the sensitivity defined above and assumed a false positive detection rate of 0. We then estimated 90% confidence intervals using a normal sampling distribution.

### Cytokine co-expression modules

Cytokine co-expression modules were identified separately for CD19-3T3 stimulated samples and anti-CD3/CD28-beads stimulated samples. We computed Pearson correlation on the concatenated normalized count matrices, removing any cytokines expressed in fewer than 10 cells and removing any associations with a Benjamini-Hochberg adjusted *q*-value greater than 0.05. The resulting correlation matrix was then clustered using hierarchical clustering. For each of the identified cytokine co-expression modules, AUCell was used to identify cells expressing that module (*45*). AUCell uses the “Area Under the Curve” (AUC) to calculate whether a module of genes is enriched within the expressed genes for each cell. First, for each module a continuous AUC score is computed based upon the rank expression of the genes in the module. From these continuous scores, a Boolean assignment to each of these modules was determined for every cell by examining the distribution of AUC scores across all the cells and setting a threshold above which the module is considered “on”. To build integrated visual representations of cells across experiments based upon the activity of these cytokine co-expression modules, the continuous AUC vectors representing the relative levels of module activity for each cell were reduced to 2 dimensions using UMAP. For each of the identified cytokine co-expression modules, Wilcoxon rank sum tests were performed between CR, RL and NR patients to determine statistically significant differences in the proportions of cells expressing that module. We only considered CAR+ cells in these tests.

### Differential Expression Analysis

All differentially expression analyses was performed using Seurat v3 (*46*). The function “FindMarkers” was used for pairwise comparison between groups of cells (samples or clusters). A log fold-change threshold of 0.25 was applied in later steps to select genes as differentially expressed. Ingenuity Pathway Analysis (IPA, QIAGEN)(*30*) was used to pathway enrichment analysis based on the differentially expressed genes associated with each cluster. Ingenuity knowledge base (genes only) was used as reference set to perform Core Expression Analysis. T cell related signaling pathways were selected from the top identified pathways to represent major functional profiles of each cluster. The z-score was used to determine activation or inhibition level of specific pathways.

### Ingenuity Pathway Analysis

Ingenuity Pathway Analysis (IPA, QIAGEN) was used to understand underline signaling pathways(*30*). Here, DEGs distinguishing each identified cluster and the corresponding fold change value of each gene were loaded into the dataset. Ingenuity knowledge base (genes only) was used as reference set to perform Core Expression Analysis. T cell related signaling were selected from identified top canonical pathways to represent major functional profiles of each cluster. The z-score was used to determine activation or inhibition level of specific pathways.

### Statistics

Analyses were performed with GraphPad Prism 8. Mann Whitney test was used to compare the expression level of specific genes or ADTs and the cell proportion of clusters. A P value<0.05 was considered statistically significant.

## Acknowledgement

The research was supported by Stand-Up-to-Cancer (SU2C) Convergence 2.0 Grant (to R.F.) and the Packard Fellowship for Science and Engineering (to R.F.). This material is based upon work supported under a collaboration by Stand Up To Cancer, a program of the Entertainment Industry Foundation and the Society for Immunotherapy of Cancer. The microfluidic devices were fabricated in the Yale School of Engineering and Applied Science cleanroom and Yale West Campus cleanroom. The sequencing service was conducted at Yale Stem Cell Center Genomics Core facility which was supported by the Connecticut Regenerative Medicine Research Fund and the Li Ka Shing Foundation or Yale Center for Genomics Analysis (YCGA). Computational data analysis was conducted with the Yale High Performance Computing clusters (HPC).

