## Supplementary Information for "Single-cell antigen-specific activation landscape of CAR T infusion product identifies determinants of CD19 positive relapse in patients with ALL"

Correspondence to:

### Contents

**Supplementary Table 1.** Patient demographics and response documentation.
**Supplementary Table 2.** Differentially expressed genes between proliferating, partially active, and fully active cell populations across the 4 experimental conditions. **Attached as separate files**.

**Supplementary Figure 1**. Flow cytometry evaluation of engineered 3T3 cells.
**Supplementary Figure 2.** Canonical signaling pathways of each cluster identified in the global clustering analysis of all the CAR T cells profiled in our project.
**Supplementary Figure 3.** Differentially expressed gene analysis of stimulation condition-related clusters.
**Supplementary Figure 4.** Detection of CAR structure, CAR gene expression and CD4/CD8 cellular molecule expression.
**Supplementary Figure 5.** Differentially expressed genes between unstimulated baseline CAR T cells from CR and RL patients and the corresponding pathways.
**Supplementary Figure 6.** Differentially expressed genes of each cluster identified in isolated CAR(+) cells from CR and RL patients.
**Supplementary Figure 7.** A four-marker scheme to determine T cell differentiation states.
**Supplementary Figure 8.** Expression of activation-related surface protein ADTs and encoding genes in CD19 stimulated CAR T cells and their comparison between response groups.
**Supplementary Figure 9.** Expression of co-inhibitory surface protein ADTs and encoding genes in CD19 stimulated CAR T cells and their comparison between response groups.
**Supplementary Figure 10.** Differentially expressed genes of each cluster identified in activated CAR(+) cells from all the patients.
**Supplementary Figure 11.** Cytokine gene expression pattern of TCR-stimulated CAR-T cells.
**Supplementary Figure 12.** Global pathway analysis and comparison of CD19-specific or anti-CD3/CD28-beads stimulated pre-infusion CAR T cells.

**Supplementary Table 1.** Patient demographics and response documentation

| ID | Infusion date | Age | Sex | Response to CART19 | B cell aplasia (BCA) | BCA duration (months) | Relapse | Time to relapse (days) | Relapse type | Last contact days |
|---|---|---|---|---|---|---|---|---|---|---|
| 112 | 4/16/13 | 9.6 | F | MRD neg CR | Yes | 72 | No | N/A | N/A | 2171 |
| 139 | 6/11/14 | 15.2 | F | MRD neg CR | Yes | 4 | Yes | 1807 | CD19 negative | 2000 |
| 151 | 9/9/14 | 15.5 | F | MRD neg CR | Yes | 61 | No | N/A | N/A | 1833 |
| 158 | 2/25/15 | 4.9 | F | MRD neg CR | Yes | 4 | No | N/A | N/A | 1696 |
| 165 | 6/23/15 | 14.5 | M | MRD neg CR | Yes | 54 | No | N/A | N/A | 1637 |
| 157 | 2/10/15 | 12.3 | M | MRD neg CR | Yes | 6 | Yes | 632 | CD19 positive | 1135 |
| 161 | 4/7/15 | 9.7 | F | MRD neg CR | Yes | 3 | Yes | 287 | CD19 positive | 602 |
| 171 | 9/8/15 | 9.8 | M | MRD neg CR | Yes | 3 | Yes | 595 | CD19 positive | 944 |
| 110 | 3/19/13 | 10.0 | F | MRD neg CR | Yes | 3 | Yes | 79 | CD19 positive | 221 |
| 111 | 4/1/13 | 9.9 | F | MRD positive | Yes | 1 | Yes | 44 | CD19 positive | 69 |
| 117 | 5/9/13 | 16.2 | M | No response | N/A | N/A | N/A | N/A | N/A | 79 |
| 167 | 7/9/15 | 21.5 | M | No response | N/A | N/A | N/A | N/A | N/A | 31 |

**Supplementary Table 2.** Differentially expressed genes between proliferating, partially active, and fully active cell populations across the 4 experimental conditions. **Attached as separate files**.

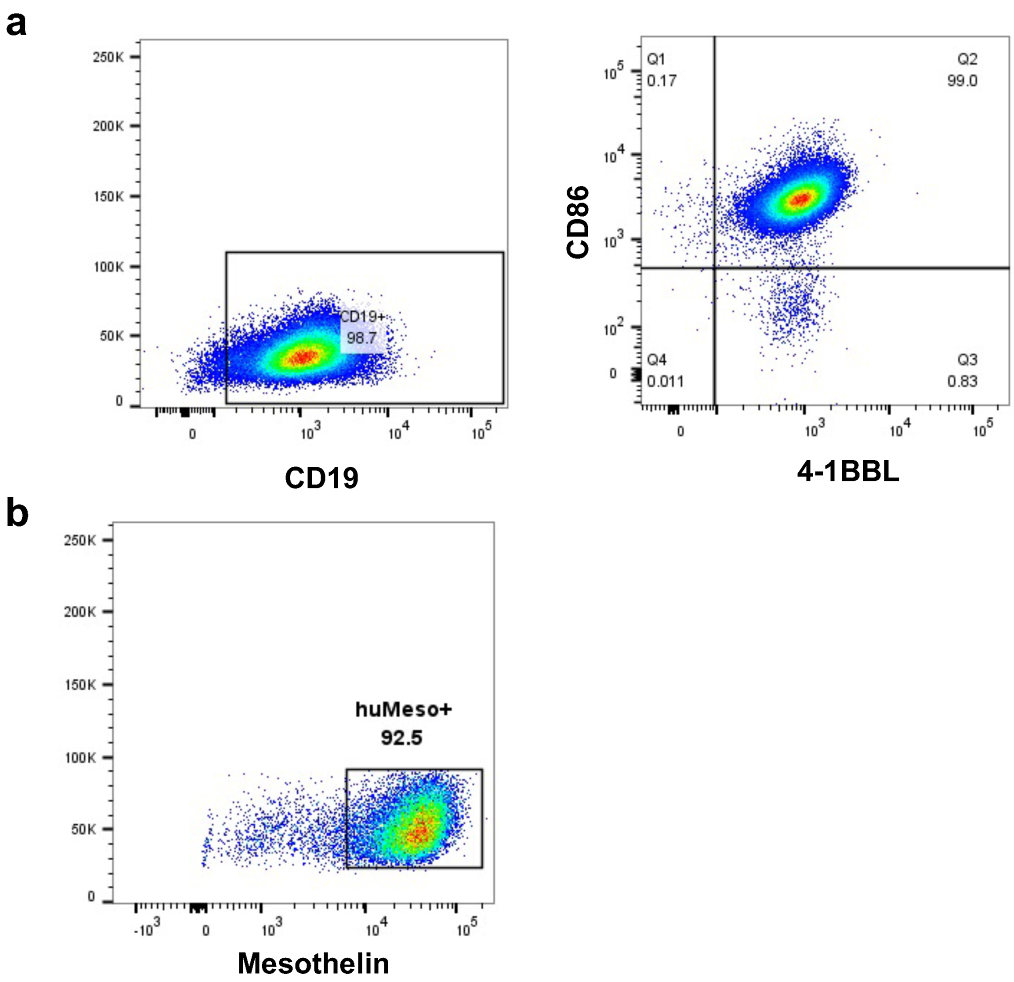


**Supplementary Figure 1. Flow cytometry evaluation of engineered 3T3 cells.** (**a**, **b**) Representative flow cytometry plots showing stable expression of human CD19, CD86, 4-1BB ligand (**a**) and mesothelin (**b**) in 3T3 cells.

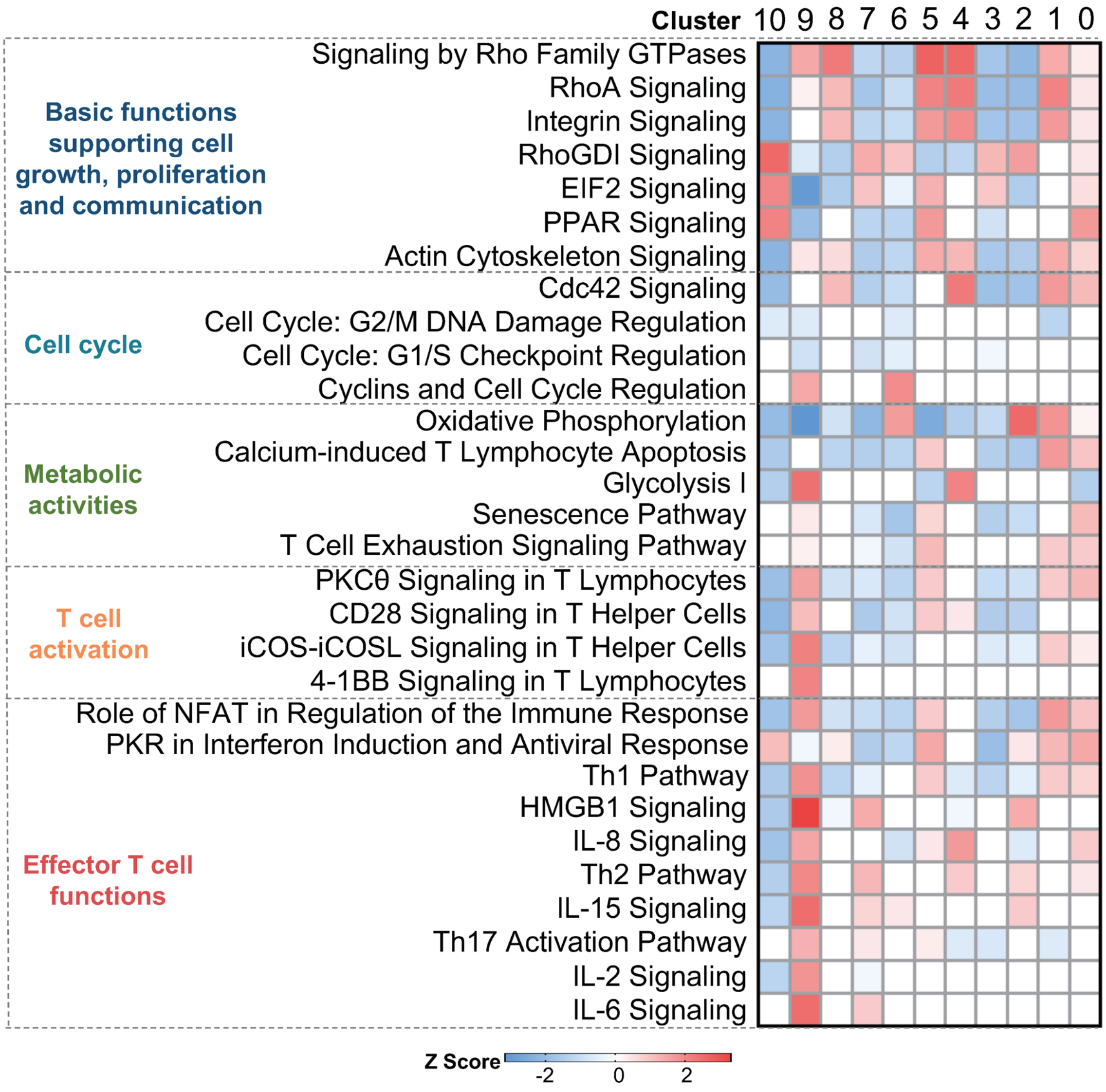


**Supplementary Figure 2. Canonical signaling pathways of each cluster identified in the global clustering analysis of all the CAR T cells profiled in our project.** Differentially expressed genes of each cluster are used to identify the biological pathways. A statistical quantity, called $z$ score, is computed and used to characterize the activation level. $z$ score reflects the predicted activation level ($z < 0$, inhibited; $z > 0$, activated; $z \geq 2$ or $z \leq -2$ can be considered significant).

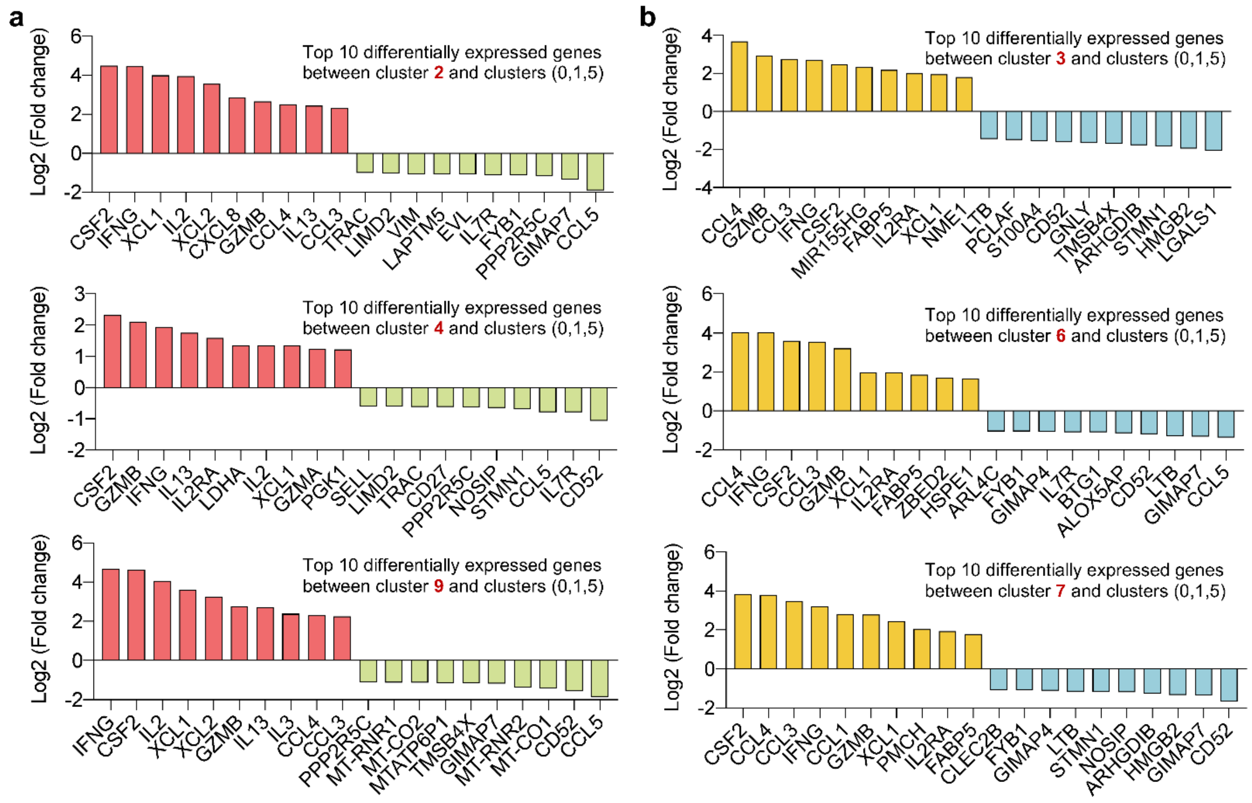


**Supplementary Figure 3. Differentially expressed gene analysis of stimulation condition-related clusters.** (**a**) Top 10 differentially expressed genes between clusters enriched in CD19 stimulated CAR T cells vs. MSLN-3T3 stimulated cells (**b**) Top 10 differentially expressed genes between clusters enriched in anti-CD3/CD28 beads stimulated CAR T cells vs. unstimulated cells.

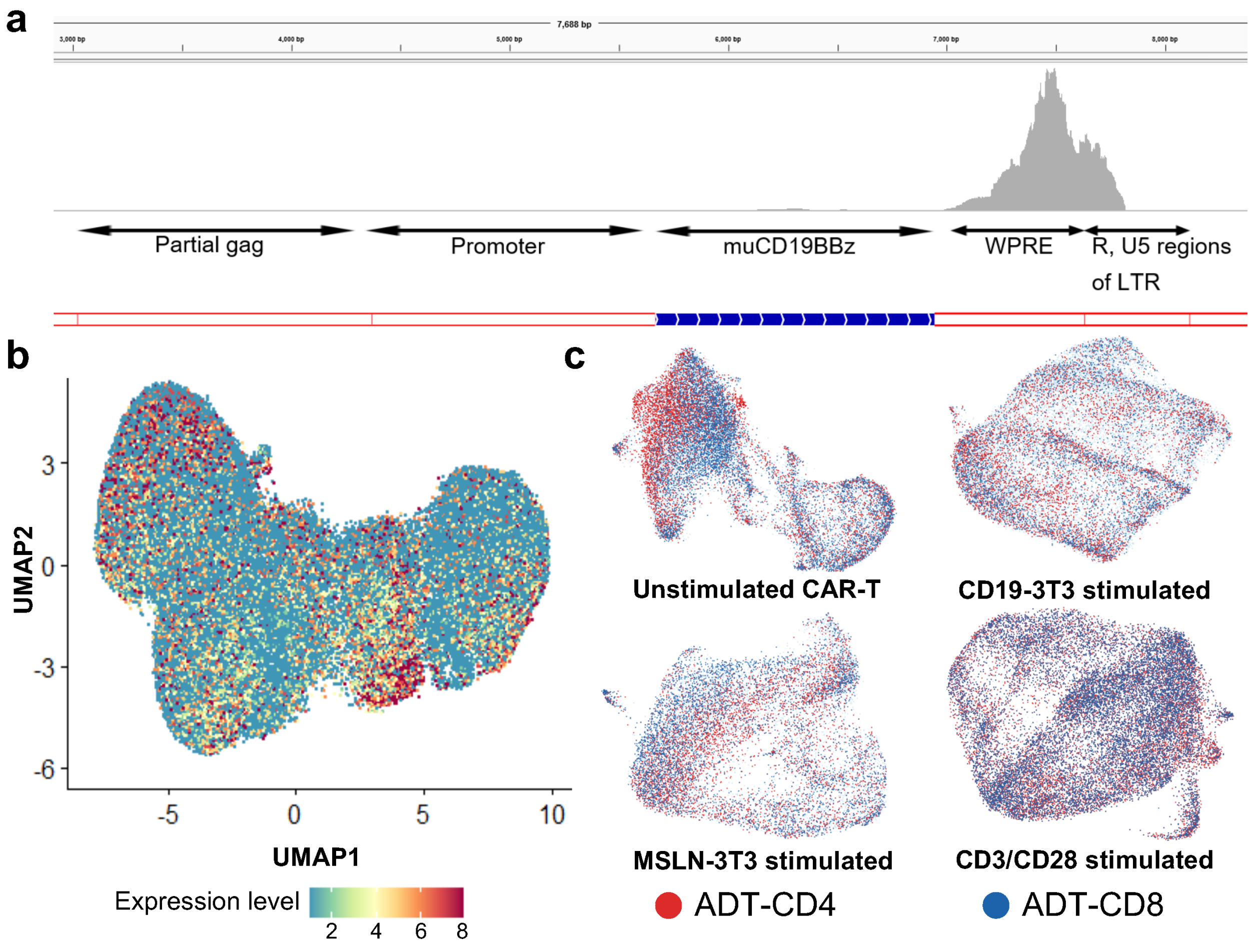


**Supplementary Figure 4. Detection of CAR structure, CAR gene expression and CD4/CD8 cellular molecule expression.** (**a**) Identify the expression of the lentiviral vector elements in the CAR construct. (**b**) The distribution of CAR gene expression among all profiled single cells. (**c**) ADT-CD4 and ADT-CD8 expression determined by CITE-seq data in four stimulation conditions.

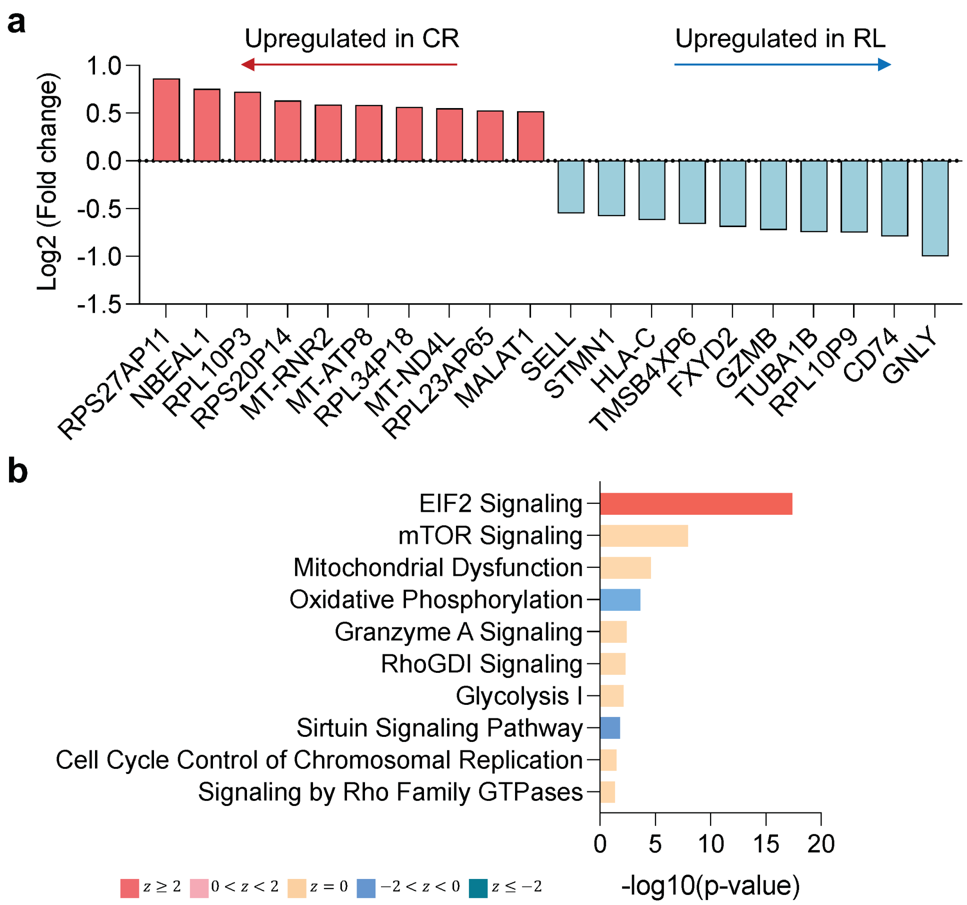


**Supplementary Figure 5. Differentially expressed genes between unstimulated baseline CAR T cells from CR and RL patients and the corresponding pathways.** (**a**) Top 10 DEGs upregulated in CR or RL group. (**b**) Corresponding canonical pathways regulated by the highly differential genes identified in (**a**). $z$ score reflects the predicted activation level ($z < 0$, inhibited; $z > 0$, activated; $z \geq 2$ or $z \leq -2$ can be considered significant).

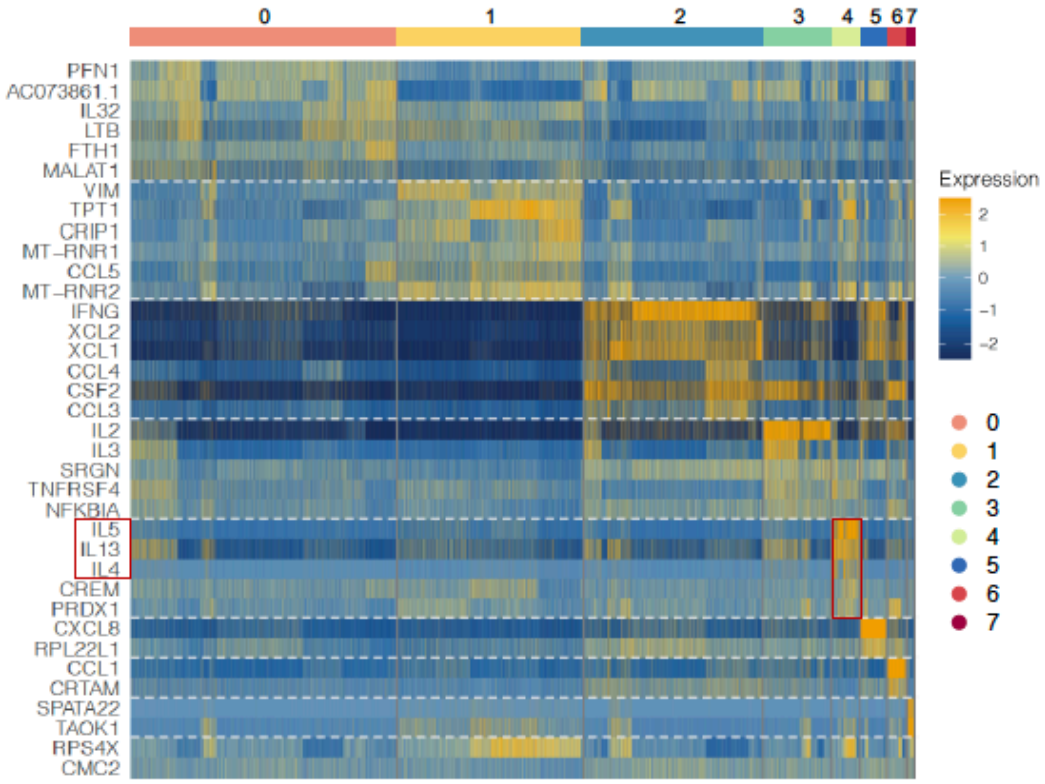


**Supplementary Figure 6. Differentially expressed genes of each cluster identified in isolated CAR(+) cells from CR and RL patients.** Heatmap of top 6 DEGs of each cluster was shown, among which the highly expression of Th2-related genes *IL13*, *IL5* and *IL4* in cluster 4 was indicated.

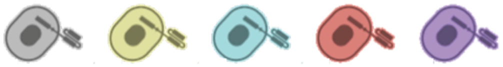

| | $T_N$ | $T_{SCM}$ | $T_{CM}$ | $T_{EM}$ | $T_{EF}$ |
|---|---|---|---|---|---|
| CD62L | + | + | + | − | − |
| CCR7 | + | + | + | − | − |
| CD45RA | + | + | − | − | + |
| CD45RO | − | + | + | + | − |

**Supplementary Figure 7. A four-marker scheme to determine T cell differentiation states.** $T_N$, naïve; $T_{SCM}$, stem cell-like memory; $T_{CM}$, central memory; $T_{EM}$, effector memory; $T_{EF}$, effector T cells.

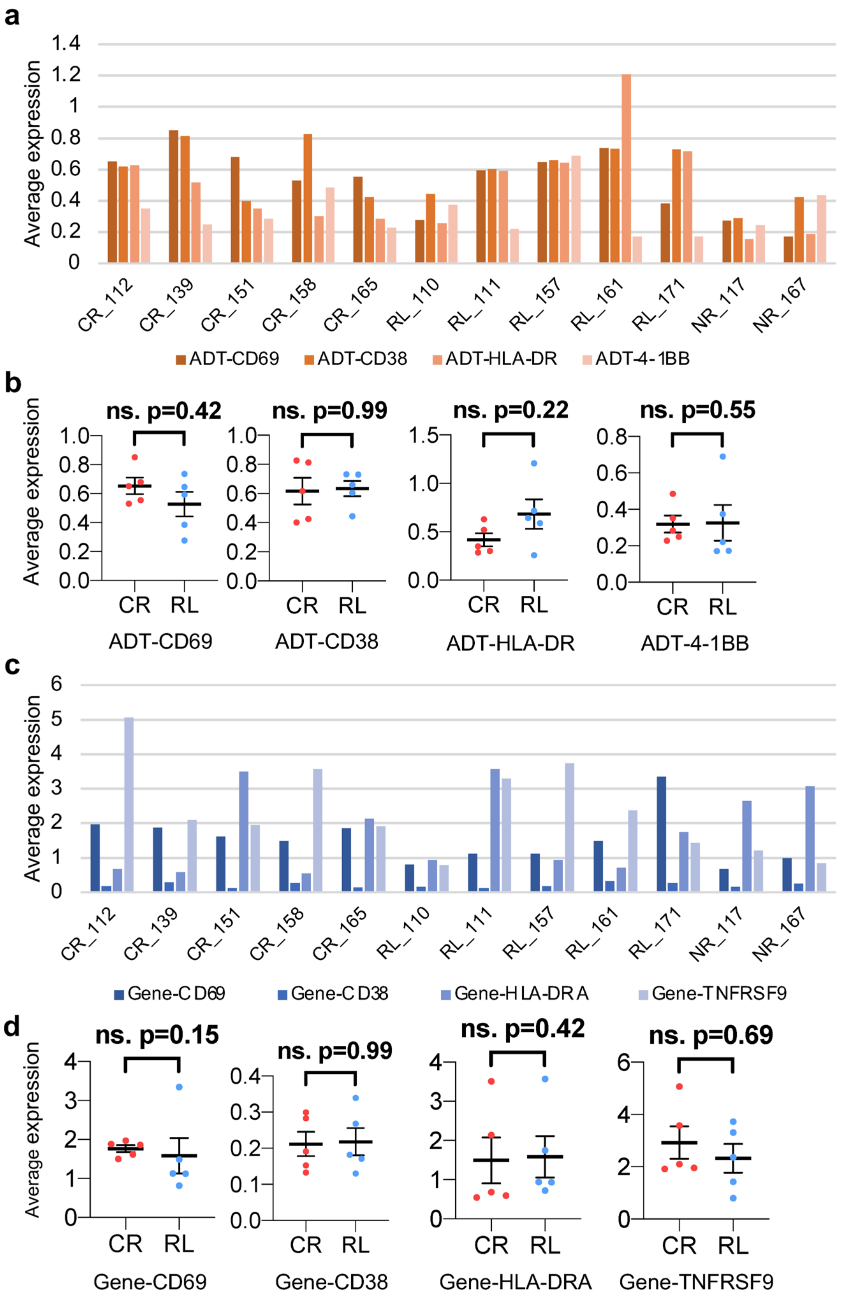


**Supplementary Figure 8. Expression of activation-related surface protein ADTs and encoding genes in CD19 stimulated CAR T cells and their comparison between response groups.** (**a**) The expression level of ADT-CD69, ADT-CD38, ADT-HLA-DR, ADT-4-1BB in each patient. (**b**) Comparison of the four protein markers expression between CR and RL patients. (**c**) The expression level of genes *CD69*, *CD38*, *HLA-DRA*, *TNFRSF9* in each patient. (**d**) Comparison of the four encoding genes expression between CR and RL patients. Each scatter point represents the average expression value of all single cells of specific patient. The $P$ values were calculated with two-tailed Mann-Whitney test. ns. Not significant. Scatter plots show mean±s.e.m.

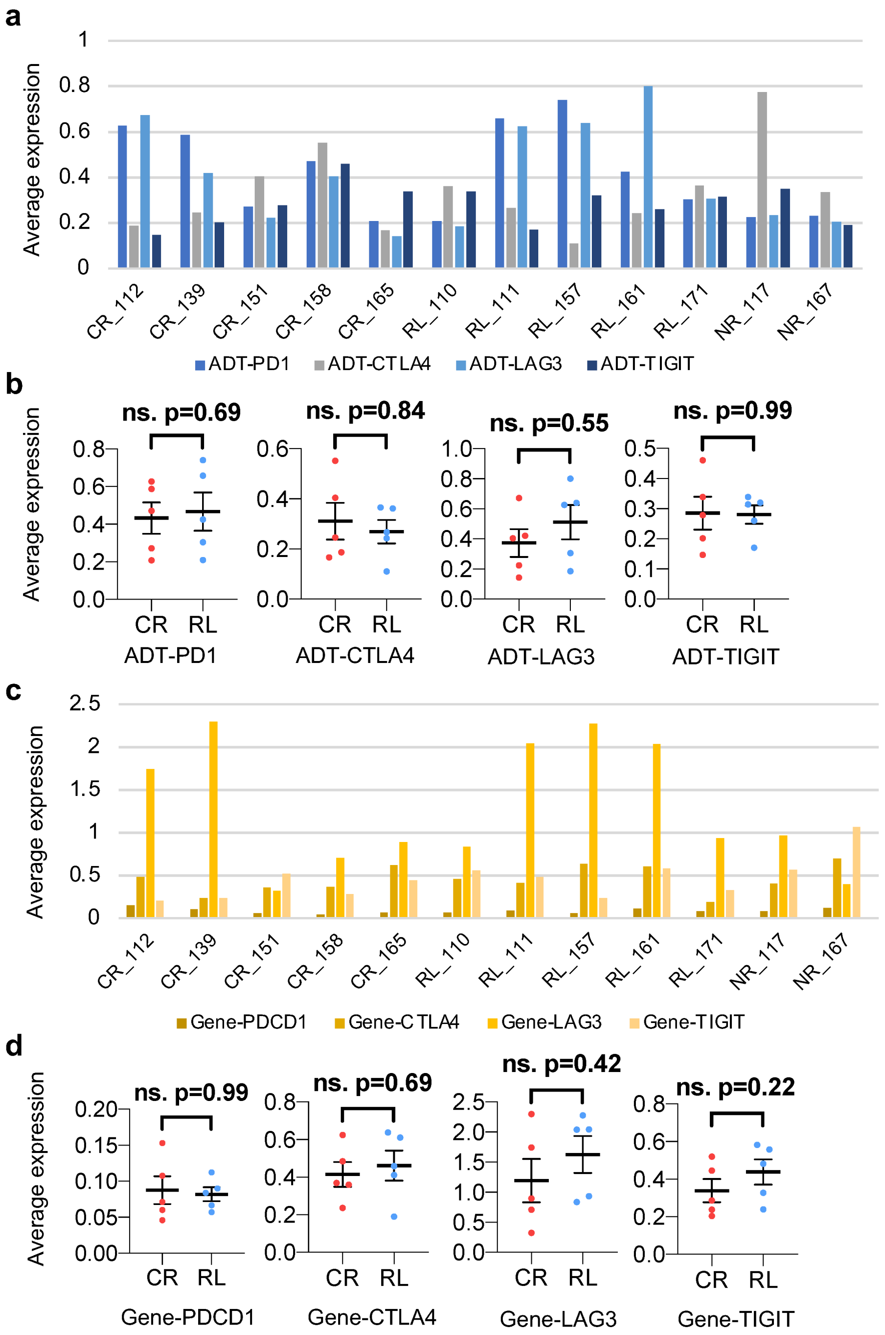


**Supplementary Figure 9. Expression of co-inhibitory surface protein ADTs and encoding genes in CD19 stimulated CAR T cells and their comparison between response groups.** (**a**) The expression level of ADT-PD1, ADT-CTLA4, ADT-LAG3, ADT-TIGIT in each patient. (**b**) Comparison of co-inhibitory protein markers expression between CR and RL patients. (**c**) The expression level of genes *PDCD1*, *CTLA4*, *LAG3*, *TIGIT* in each patient. (**d**) Comparison of co-inhibitory genes expression between CR and RL patients. Each scatter point represents the average expression value of all single cells of specific patient. The *P* values were calculated with two-tailed Mann-Whitney test. ns. Not significant. Scatter plots show mean±s.e.m.

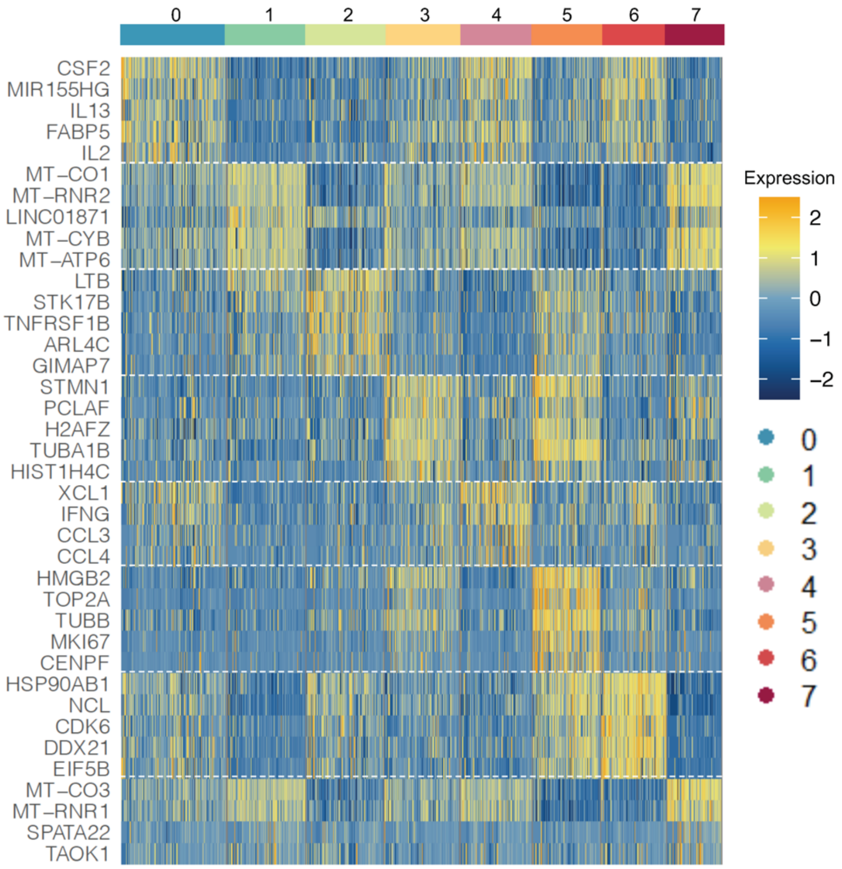


**Supplementary Figure 10. Differentially expressed genes of each cluster identified in activated CAR(+) cells from all the patients.** Heatmap of top 5 DEGs of each cluster was shown.

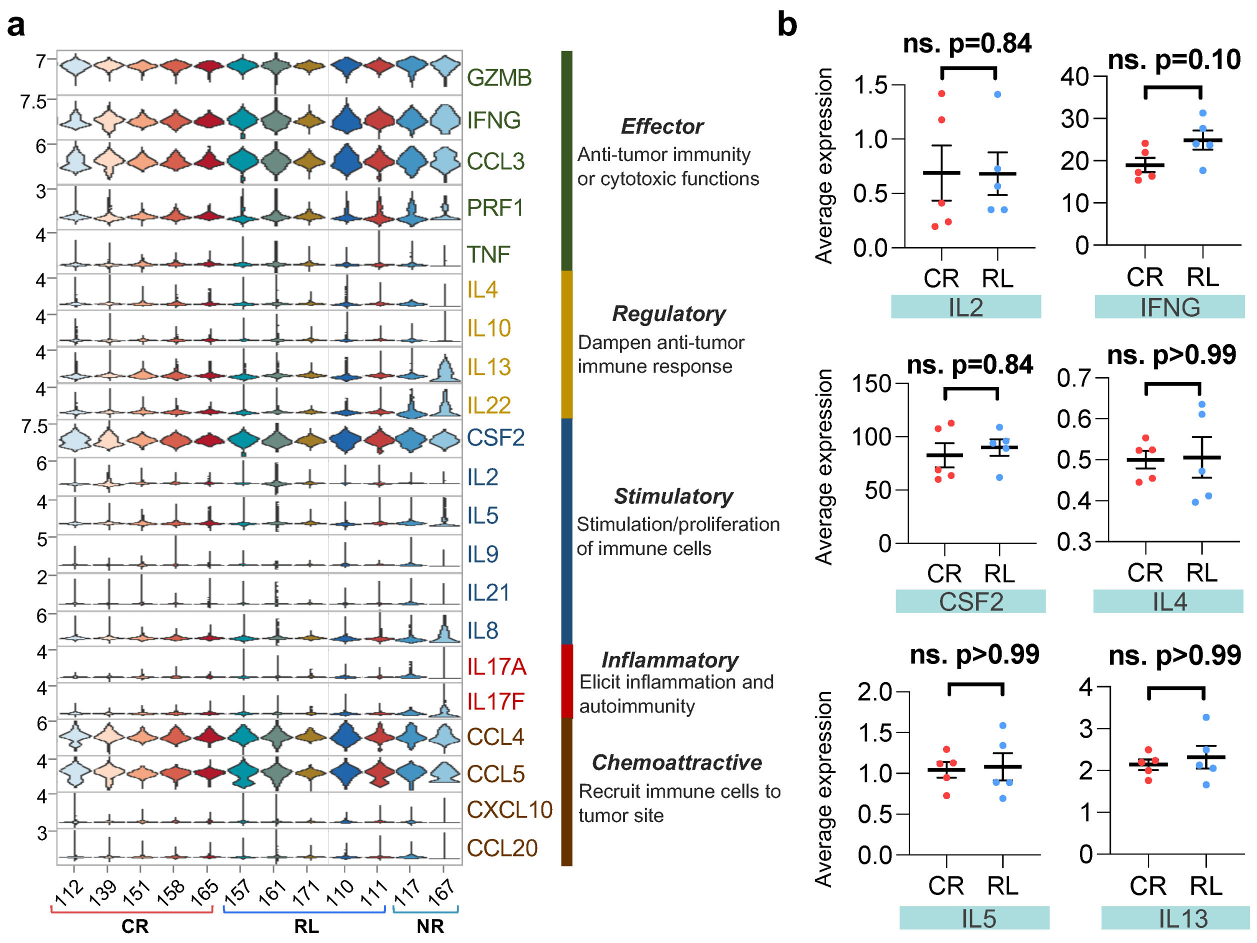


**Supplementary Figure 11. Cytokine gene expression pattern of TCR-stimulated CAR-T cells.** (**a**) Single cell expression level violin plot of all key immunologically relevant cytokine genes from CAR(+) cells. (**b**) Quantification of expression level of selected cytokine genes and comparisons between CR and RL group. Each scatter point represents the average expression value of all single cells of specific patient. The $P$ values were calculated with two-tailed Mann-Whitney test. ns. Not significant. Scatter plots show mean±s.e.m.

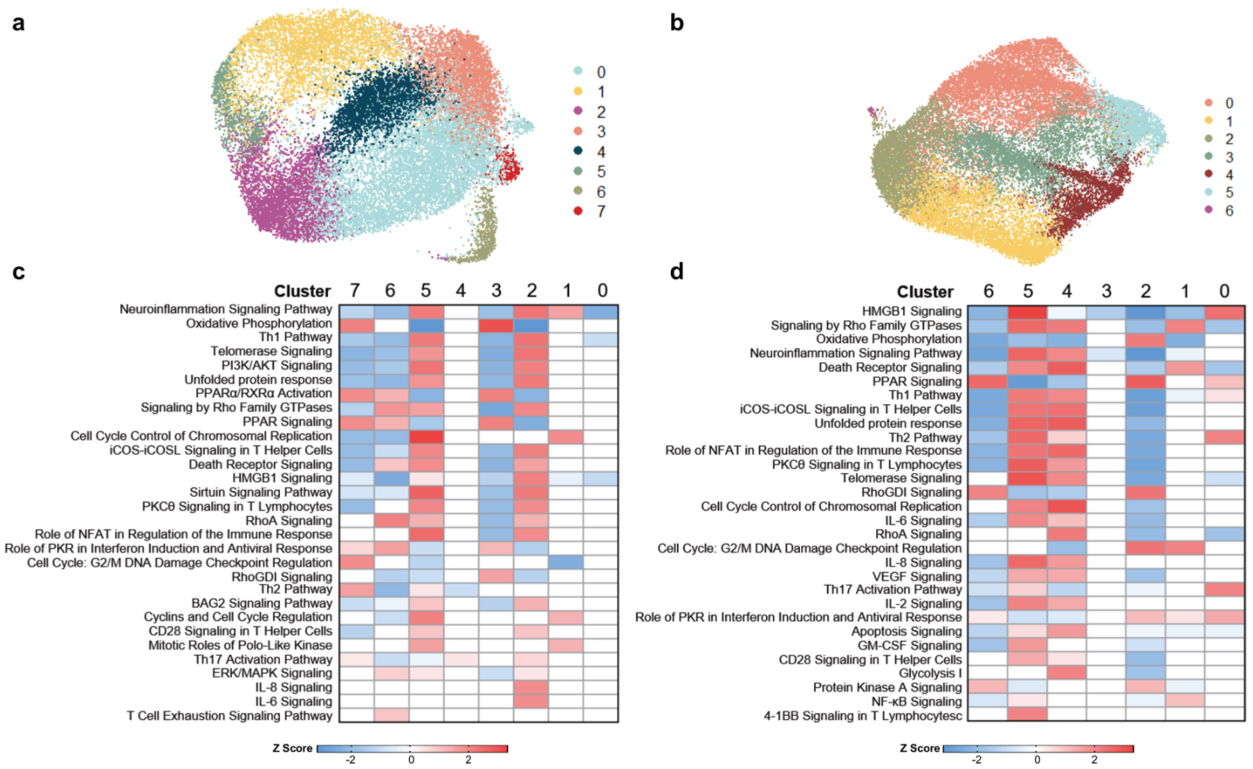


**Supplementary Figure 12. Global pathway analysis and comparison of CD19-specific or anti-CD3/CD28-beads stimulated pre-infusion CAR T cells.** (**a**) UMAP clustering of anti-CD3/CD28 beads stimulated cells. (**b**) UMAP clustering of CD19-3T3 stimulated cells. (**c**) Canonical pathway analysis and comparison of clusters in (**a**). (**d**) Canonical pathway analysis and comparison of clusters in (**b**). Differentially expressed genes of each cluster are used to identify the biological pathways. $z$ score reflects the predicted activation level ($z < 0$, inhibited; $z > 0$, activated; $z \geq 2$ or $z \leq -2$ can be considered significant).
